# Attention Deficit Hyperactivity Disorder and the gut microbiome: An ecological perspective

**DOI:** 10.1101/2022.08.17.504352

**Authors:** Trevor Cickovski, Kalai Mathee, Gloria Aguirre, Gorakh Tatke, Alejandro Hermida, Giri Narasimhan, Melanie Stollstorff

## Abstract

Attention Deficit Hyperactivity Disorder (ADHD) is an increasingly prevalent neuropsychiatric disorder characterized by hyperactivity, inattention, and impulsivity. Symptoms emerge from underlying deficiencies in neurocircuitry, and recent research has suggested a role played by the gut microbiome. The gut microbiome is a complex ecosystem of interdependent taxa with an exponentially complex web of interactions involving these taxa, plus host gene and reaction pathways, some of which involve neurotransmitters with roles in ADHD neurocircuitry. Studies have analyzed the ADHD gut microbiome using macroscale metrics such as diversity and composition, and have proposed several biomarkers. Few studies have delved into the complex underlying dynamics ultimately responsible for the emergence of such metrics, leaving a largely incomplete, sometimes contradictory, and ultimately inconclusive picture.

We aim to help complete this picture by venturing beyond taxa *abundances* and into taxa *relationships* (i.e. cooperation and competition), using a publicly available gut microbiome dataset from 30 Control (15 female, 15 male) and 28 ADHD (15 female, 13 male) undergraduate students. We conduct our study in two parts. We first perform the same macroscale analyses prevalent in ADHD gut microbiome literature (diversity, differential, biomarker, and composition) to observe the degree of correspondence, or any new trends. We then estimate two-way ecological relationships by producing Control and ADHD Microbial Co-occurrence Networks (MCNs), using SparCC correlations (*p* < 0.01). We perform community detection to find clusters of taxa estimated to mutually cooperate along with their centroids, and centrality calculations to estimate taxa most vital to overall gut ecology. We conclude by summarizing our results, and provide conjectures on how they can guide future experiments, some methods for improving our experiments, and general implications for the field.

## Introduction

Attention Deficit Hyperactivity Disorder (ADHD) is a significant mental health problem with a current 3.4% prevalence worldwide (1). In the United States, ADHD affects one in 10 children (a 43% increase over the last 15 years) (2), and 3-16% of adults (3) with that percentage increasing over the past 20 years. Individuals with ADHD face many practical challenges, including risk for low academic achievement, lower employment status, and incarceration (4). Symptoms of hyperactivity, impulsivity, and inattention characterize ADHD (5). Underlying ADHD behavioral symptoms are deficits in the neurocognitive mechanisms of both executive function (EF) and emotional regulation (ER) (6), including and extending beyond prefrontal-striatal networks (7). EF refers to a set of cognitive control processes, includes one’s ability to focus on relevant information while suppressing irrelevant distractors. ER generally ascribes to one’s ability to effectively cope with emotionally charged circumstances (both negative and positive). Many medications have been developed to combat the disorder by influencing the underlying neurocircuitry (8).

The pathogenesis of ADHD is thought to be multifactorial, with heritability estimates at roughly 70-90% (9). These genetic connections suggest some dependency on underlying metabolic reactions, directly or indirectly involving gene products. In the meantime, the new and exciting field of microbiome research has made its way into the mental health domain. Our gut is home to a plethora of bacteria, fungi, and other microbial organisms, whose collective genomes comprise our gut *microbiome*. Studies estimate that the average number of bacterial cells in humans matches or exceeds that of host cells (10,11). Each bacterium has unique genetic material that produces different sets of metabolites, which interact with each other and host metabolites downstream (12), creating a complex host-microbiome web of interactions. It has become increasingly important to pay attention to the symbiotic relationship between the gut microbiome and brain development and function, often referred to as the *gut-brain-microbiome axis* (13). This axis is a bidirectional communication network, providing gut microbiota and metabolites an avenue for influencing brain development and function (14–18). One proposed mechanism through which gut microbiota may affect our neurobiology is by altering the levels of *neurotransmitters*, including dopamine and serotonin (5-HT) (19), which fuel brain regions that mediate cognition and emotion. Although serotonin is also produced in the brain, up to 90% of serotonin is synthesized in the gut (20).

Connections between the gut microbiome and neurotransmitters, EF/ER, and *neuropsychiatric disorders (NPDs)* characterized by EF/ER disorders are already well-established. In rodents, anxiety and social behavior have been linked to the gut microbiome that can be attributed to altered neurotransmission in the hippocampus and amygdala (21). In humans, associations between microbiome composition and ER have been shown (18). It has also been established that the gut microbiome can release dopamine and 5-HT, impacting ER (22,23). Connections on the cognitive axis related to EF are less well-established in humans, though some theories are beginning to emerge (24). In humans, dopamine influences EF (25). In rodents, the gut microbiome is linked to dopamine (26), and EF-like behavior (27). The Autism Spectrum Disorder (ASD) (28), which is associated with impaired EF (29), has been linked to the gut microbiome (30). In animal studies, the gut microbiome has been associated with anxiety-related disorders such as depression (31–36). People with stress-related diseases have responded positively to probiotics (37,38). Connections between the gut microbiome and another neuropsychiatric disorder (NPD) characterized by EF/ER dysfunction such as ADHD would further support the impact of the gut microbiome on EF/ER. It could also help to explain the large amount of symptomatic overlap that exists between ADHD with other NPDs, particularly ASD (39–41), and could even provide differentiating factors (42) to help address the current diagnosis challenges due to this overlap (43), and new potential options for treatment (44). The fact that individuals with ADHD suffer from gastrointestinal (GI) dysfunction, including childhood digestive difficulties and low-grade inflammation (45) as well as constipation (46,47), only further suggests a potential role of the gut microbiome in this disorder.

There are limited studies that implicate the gut microbiome on clinically diagnosed ADHD, and recent efforts have been made to survey and summarize their results (48–51). Two in particular published this year (49,50) contained findings from every published study involving ADHD and the gut microbiome. Based on this, we make the following observations about the current state of ADHD and gut microbiome research:

1. ***Diversity results are <u>contradictory</u> and <u>inconclusive</u>***. Even with closely age-matched gut microbiome studies using the same Shannon index (52) to measure alpha-diversity, one set (mean age 11.9 years) revealed a lower level of alpha-diversity in ADHD patients (53), another (mean age 9.3 years) revealed higher alpha-diversity (54), a third (ages 6-10) reported no difference at all (55), and a fourth (10- and 15-year-olds) (56) reported higher alpha-diversity in ADHD 15-year-olds, but no difference in ADHD 10- year-olds. Within these same four studies, the first (53) reported a beta-diversity difference between ADHD and Control, while the other three reported no difference (54–56). With a mean age only slightly higher (20.2), a fifth study found no alpha-diversity difference, but a beta-diversity difference (57).
2. ***Many biomarkers have been proposed, some contradictory, others mixed depending on taxonomic level, and others inconclusive***. Proposed ADHD biomarkers include: increased *Collinsella* (58) (phylum Actinobacteria), increased *Fusobacterium* (54) (Fusobacteria), decreased *Lachnospiraceae* (59), *Lactobacillus* (54,60), and *Ruminococcus gnavus* (59) (all Firmicutes), decreased *Prevotella*/*Porphyromonadaceae* (53) and increased *Paraprevotella xylaniphila*, *Odoribacteriaceae* and member species *Odoribacter splanchicus* (59) (all Bacteroidetes), decreased *Haemophilius* (57) and increased *Neisseria* (53),*Sutterella stercoricanis* (54), and *Desulfovibrio* (61) (all Proteobacteria).

More mixed results have been reported with respect to the following taxa:

**Clostridiales (Firmicutes).** This order (57) was reported as increased in studies involving ADHD children and adolescents, but another study involving 18-24 month-olds (60) found members of this order as lower.

***Ruminococcaceae* (Firmicutes).** This family was reported as elevated in ADHD by one study (57) along with member genus *Ruminococcus*, but member genus *Faecalibacterium* was reported as reduced in two others (55,59).

***Veillonellaceae* (Firmicutes).** Within the same study (59), family *Veillonellaceae* and genus *Veillonella* were reduced in ADHD, but member species *V. parvula* was elevated.

***Bacteroidaceae* (Bacteroidetes).** *Bacteroidaceae* was found as elevated in ADHD by one study (53), member genus *Bacteroides* was reduced in another among 18-month-olds (60), member species *B. uniformis* (54), *B. ovatus* (54) and *B. coccae* (59) were all reported as elevated, and member species *B. coprocola* was reported as reduced (54).

For one particular taxon, results have been contradictory:

***Bifidobacterium* (Actinobacteria).** Perhaps no greater mystery currently exists than the role of genus *Bifidobacterium*. One Dutch study found a nominal increase in *Bifidobacterium* with average ADHD and Control subject ages of 19.5 and 27.1 years, respectively (62). A longitudinal study (3 months, six months and 13 years) made a somewhat contradictory observation of reduced *Bifidobacterium* during infancy, but not at age 13 (60). A third study (58) reported reduced *Bifidobacterium* (specifically *B. longum* and *B. adolescentis*) in ADHD children (mean age: 9.3) that actually reversed after micro-nutrient treatment, where elevated *Bifidobacterium* was observed at high ADHD-Rating Scale IV (ADHD-RS-IV, (63)) scores.

The current picture of the role played by the gut microbiome in ADHD is therefore still unclear. Most of the effort to connect ADHD to the gut microbiome has involved (1) macroscale population metrics such as diversity, and/or (2) taxa abundances. These properties are in reality emergent from a complex and interdependent interaction web inolving taxa, their gene products, and those of the host (64). Diversity and abundance therefore ignore many underlying details behind their measurements, helping to explain the current incomplete picture. Venturing deeper into this web is critical to completing more of this picture. Two studies have attempted this task, both using multi-omics. One (59) reported differences ADHD neurotransmitter pathways. A second (62) uncovered a connection between *Bifidobacterium* and cyclohexadienyl dehydratase (CDT) abundances.

We have thus far only scratched the surface of this large and exponentially complex interaction web, and every completed piece has value. Multi-omics will continue to be critical, bridging an important gap between taxa, products, and metabolic reactions. We aim to complete another piece, that involves *ecological relationships* between taxa. Microbial taxa have been shown to demonstrate a wide variety of ecological relationships, including cooperation (65,66) and competition (67), that ultimately impact collective functionality of the ecosystem and macroscale properties (64). We estimate these relationships for Control and ADHD datasets and report results; including relationships, communities, driver taxa (or ‘centroids’) of these communities, and taxa central to overall gut ecology. Results can offer guidance on potential taxa to target for further multi-omics or laboratory experiments. The ultimate goal is to increase depth of knowledge about connections between the influence of the gut microbiome on an NPD that impacts millions of individuals worldwide.

This work involves two parts, conducted on a publicly available, gender-matched dataset of 16S gut microbiome sequences. The first involves performing the same macroscale analyses currently prevalent in ADHD gut microbiome literature, to note how this dataset compares, as well as any new and interesting trends. Metrics will include alpha- and beta-diversity, Sparse Partial Least Squares Discriminant Analysis (sPLS-DA, (68)) to estimate Control and ADHD differentiation degree, biomarker analysis using Linear discriminant analysis Effect Size (LEfSe, (69)), and QIIME (70) normalized abundance compositional profiles.

In the second part we estimate ecological relationships (71) within Control and ADHD gut microbiomes. We first use Microbial Co-occurrence Networks (MCNs, (72)) to estimate these relationships (73), and then perform cluster analysis using the Affinity Propagation (AP, (74)) algorithm to discover communities of mutually supporting taxa, as well as driver or ‘centroid’ nodes of these communities. Finally we perform centrality analysis using the Ablatio Triadum (ATria, (75)) algorithm, to estimate taxa most significant to the overall ecosystem.

## Materials and Methods

We provide more details on the methods we use for analysis. Our entire downstream analysis pipeline has been built using Plugin-Based Microbiome Analysis (PluMA, (76)) and is available for download within its publicly available pipeline pool.

### Cohort

We start from a publicly available dataset (Accession Number: PRJNA656791) of gut microbiome samples from an undergraduate student population. Full sequencing details are provided in the BioProject description; 16S rRNA (V3-V4 region) sequencing was used, following steps corresponding to standard Illumina protocols (77). Each deidentified sample provides gender and ADHD assessment based on Adult ADHD Self Report Scale (ASRS) score (Control, ADHD Combined, ADHD Inattentive, or ADHD Hyperactivity) in its title. For both subscales they used an ASRS score of 17 as an ADHD threshold, which also follows published practices (78). The project released 58 samples: 30 Control and 28 ADHD, with 15 females in both groups. We summarize statistics in Table 1. Of the ADHD cohort, 17 were ADHD-combined (inattentive and hyperactive), five ADHD-hyperactive, and six ADHD-inattentive (Table 1). Analyzed with a *t*-distribution, we found no significant impact of gender (*p* > .2).

**Table 1.** Cohort analyses

| Table 1. Cohort analyses |  |  |  |  |  |  |
| --- | --- | --- | --- | --- | --- | --- |
| Groups | Total No | Gender | Total No | Inattentiveness | Hyperactivity | Combined |
| | | | | ASRS Scores ± SD* | | |
| Control | 30 | Female | 15 | 11.2 ± 4.3 | 9.6 ± 4.9 | 20.8 ± 7.2 |
| | | Male | 15 | 9.8 ± 3.4 | 9.9 ± 4.4 | 19.7 ± 6.8 |
| ADHD | 28 | Female | 15 | 22.6 ± 6.8 | 19.5 ± 5.3 | 42.1 ± 10.1 |
| | | Male | 13 | 18.2 ± 3.8 | 17.3 ± 4.7 | 35.5 ± 6.1 |
| * *t*-test revealed no significant impact of gender ($p > .2$). | | | | | | |

To establish an initial set of taxa we took these sequences and compiled, clustered, and analyzed them using QIIME 1.9.1 (70), (similarity threshold of 97%, GreenGenes reference database (79)). We removed all singletons and scarce taxa (present in less than 50% of the samples) for both groups to produce our final set for analysis.

### Part I: Traditional Macroscale Analyses

We first perform macroscale analyses on this ADHD dataset that have been performed on other ADHD datasets, compare and contrast our results with those in the literature, and take note of any new and interesting observations.

#### Diversity analysis

Alpha- and beta-diversity plots were constructed using QIIME (version 1.9.1), with default metrics: observed_species (unique taxa count), Chao1 (80), and PD_whole_tree (phylogenetic diversity), and default parameters.

#### Discriminant analysis

Our study uses Sparse Partial Least Squares Discriminant Analysis (sPLS-DA, (68)), a sparse version of the Partial Least Squares (PLS, (81)) method, as a supervised method for determining differentiation degree with respect to taxa relative abundance (82).

#### Biomarker analysis

Our biomarker analysis used the <u>L</u>inear discriminant analysis <u>Ef</u>fect <u>S</u>iz<u>e</u> (LEfSe,(69)) algorithm (*p* < 0.05, LDA effect size > 2).

#### Compositional analysis

We use QIIME 1.9.1 (70) to generate compositional bar graphs, producing one bar per sample broken down by taxa percentages.

### Part II: Ecological Relationships

#### Co-occurrence network analysis

We computed correlations based on taxa relative abundances using SparCC (83) (*p* < 0.01), and built Microbial Co-occurrence Networks (MCNs, (72)) using taxa as nodes and correlations as edges. MCNs were visualized using Cytoscape (84) with layout produced by Fruchterman-Reingold (85).

#### Clustering

MCNs were clustered using Affinity Propagation (AP, (86)). AP has been shown to operate efficiently and successfully on signed and weighted biological networks without requiring an initial cluster count estimate, and additionally computes the most representative or *centroid* node for each cluster.

#### Centrality analysis

We use <u>A</u>blatio <u>Tria</u>dium (ATria, (75)) for evaluating the importance, or *centrality*, of taxa in our MCNs. ATria computes centrality for signed and weighted networks through a modified economic payment model (87) that calculates the influence of a node on all other nodes. ATria provides an alternative perspective by considering relationships (not relative abundance) when computing centrality, and unlike biomarker analysis does not compare sample sets. ATria produces a ranked list of important taxa and runs iteratively; once a taxon is found as central, ATria removes this taxon and its dependencies using social network theory (88). Then it runs again to produce the next most important taxon, repeating until no edges are left. Taxa not found as important are simply not ranked.

We analyze these ecological relationship <u>at all taxonomic levels</u> starting from phylum. We first observe the upper three levels (phylum, class, and order) for an overview of relationships between consistently abundant taxa. We then move to the lower three levels (family, genus and species) which provide a finer level of granularity and enough taxa to perform meaningful community analyses.

## Results

### Part I. Traditional Macroscale Analyses

#### Diversity

QIIME (70) alpha- and beta-diversity results produced no conclusive differences between ADHD and Control (Fig. 1). Although Fig. 1A shows a marginal Alpha diversity increase for ADHD using all three metrics: observed_species (count of unique taxa), Chao1 (80), and PD_whole_tree (phylogenetic diversity), error bars clearly indicate inconclusive results. Beta-diversity with unweighted and weighted Unifrac (89) distance also shows no separation (Fig. 1B). This lack of alpha- and beta-diversity differences matches several results from other datasets (55,56,62).

**Fig. 1.**
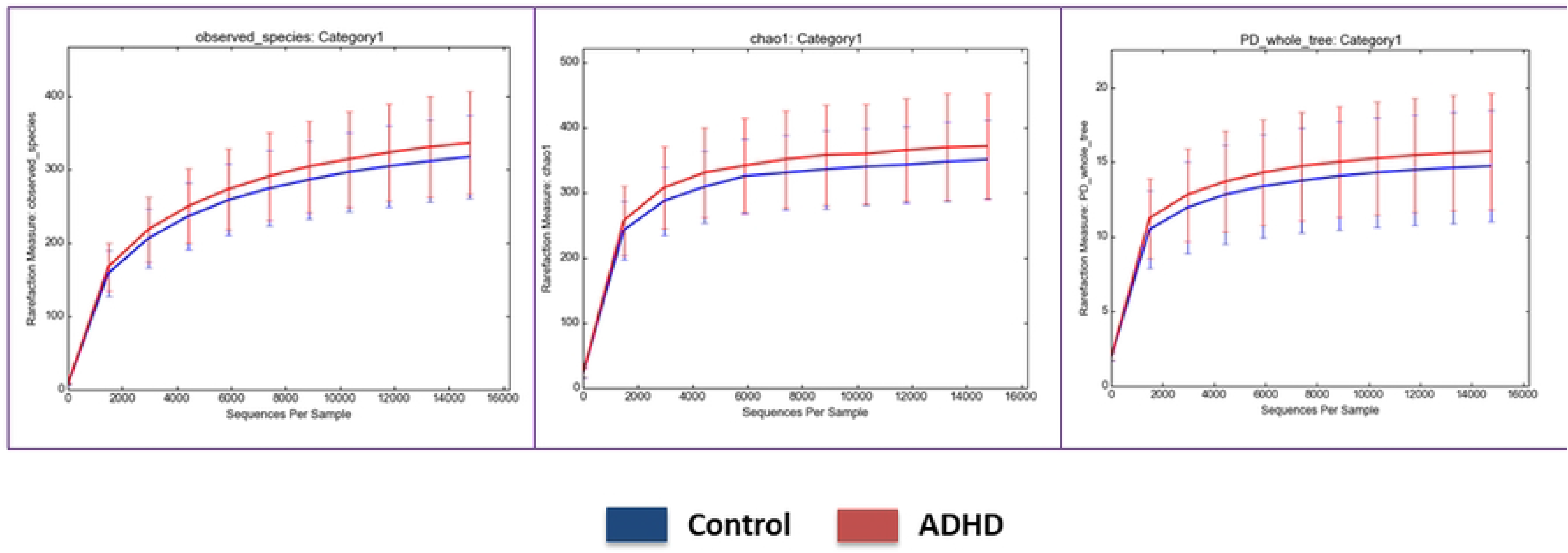
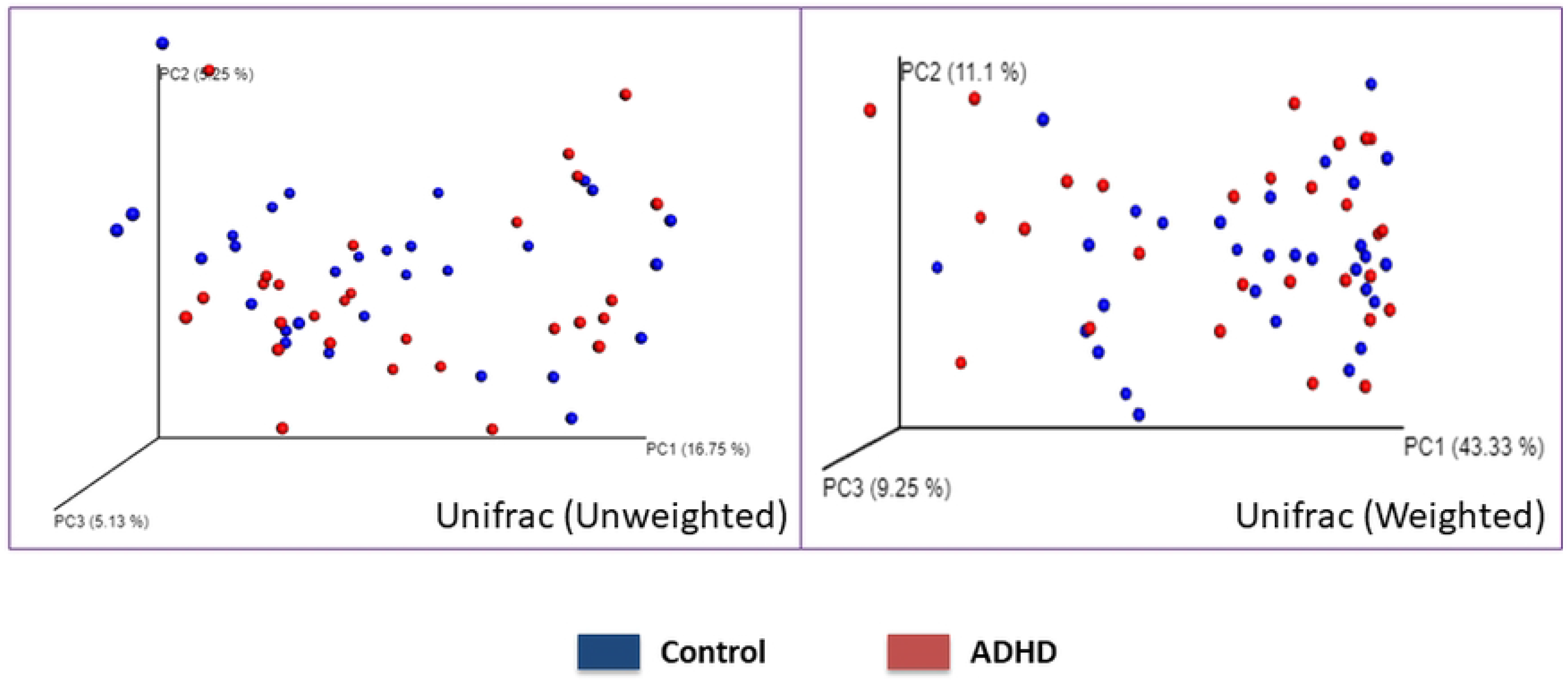
Alpha- and Beta-Diversity. (a) Alpha-diversity of Control and ADHD samples using (in order) the count of unique taxa, Chao1 richness (80), and phylogenetic diversity. (b) Beta-diversity of Control and ADHD samples computed using unweighted and weighted Unifrac (89) distance.

#### Discriminant

Discriminant analysis determines differentiation degree between datasets, accounting for all variables in each set (90). Unsupervised and supervised approaches can be used, with supervised having prior sample classification knowledge (i.e., Control or ADHD). One ADHD gut microbiome study (53) attempted the unsupervised method non-parametric multi-dimensional scaling (NMDS, (91)), but could not differentiate the two groups. Limited studies have further decomposed ADHD samples by subscale but these focus on diversity and composition, noting inattention (elevated *Dialister* and reduced *Phascolarctobacterium* (57)) and hyperactivity (lower alpha-diversity and elevated *Parabacteroides* (53)) properties.

We attempt the supervised Sparse Partial Least Square Differential Analysis (sPLS-DA, (68)), with taxa relative abundances as variables. Fig. 2(a)-(b) (ellipse confidence level=95%) shows even a supervised method cannot differentiate the groups, in general or by subscale. This is significant, as supervised approaches like sPLS-DA have *a priori* sample category knowledge and can sometimes differentiate completely random data (82). sPLS-DA did differentiate the two sets with scarce taxa present, showing some separation between Control, ADHD samples high on one subscale, and ADHD samples high on both (Fig. S1). However, a supervised method differentiating the sets only when scarce taxa (present less than half of the time) are counted shows very little.

**Fig. 2.**
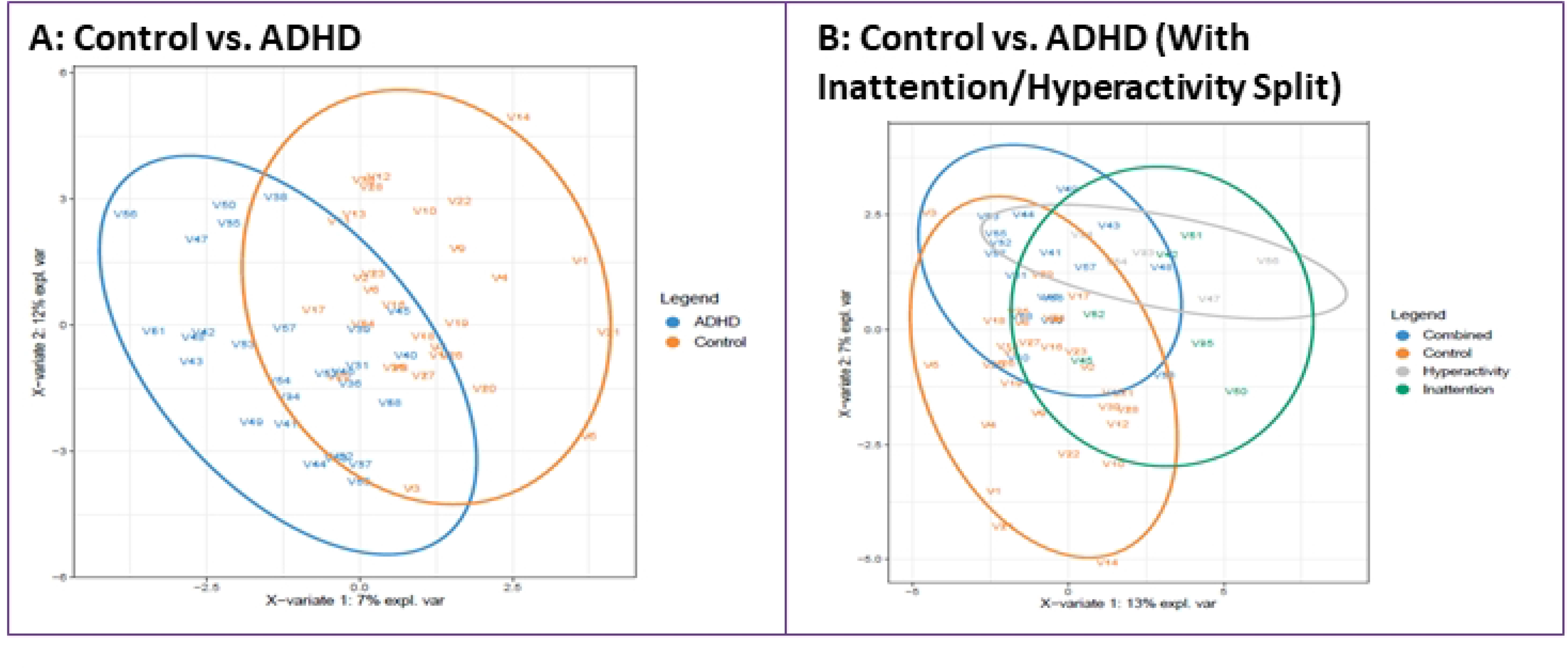
Discriminant Analysis. Results of running sPLS-DA (68) on microbiome abundance data (ellipse confidence level 95%). The figures show the analyses (a) comparing Control (orange) and ADHD (blue) groups and (b) further separating the ADHD group into inattention (green), hyperactive (grey), and combined (blue).

#### Biomarker

When performing LEfSe (69), we returned to a single ADHD set (no subscale split) to ensure roughly level sample counts with Control. Results are shown both as a cladogram (Fig. 3A) and a bar graph (Fig. 3B). LEfSe has identified orange taxa as Control biomarkers, and purple taxa as ADHD biomarkers.

**Fig. 3.**
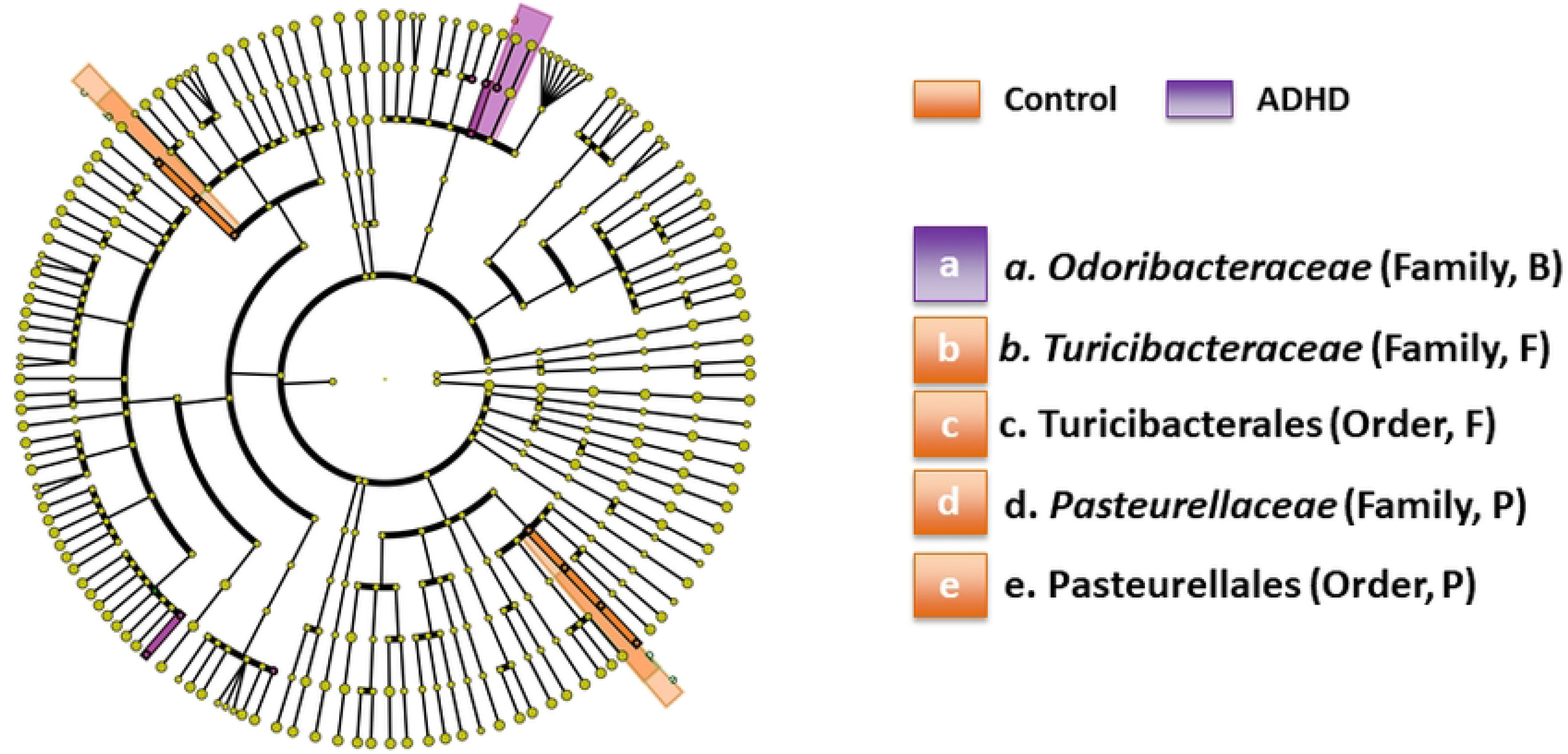
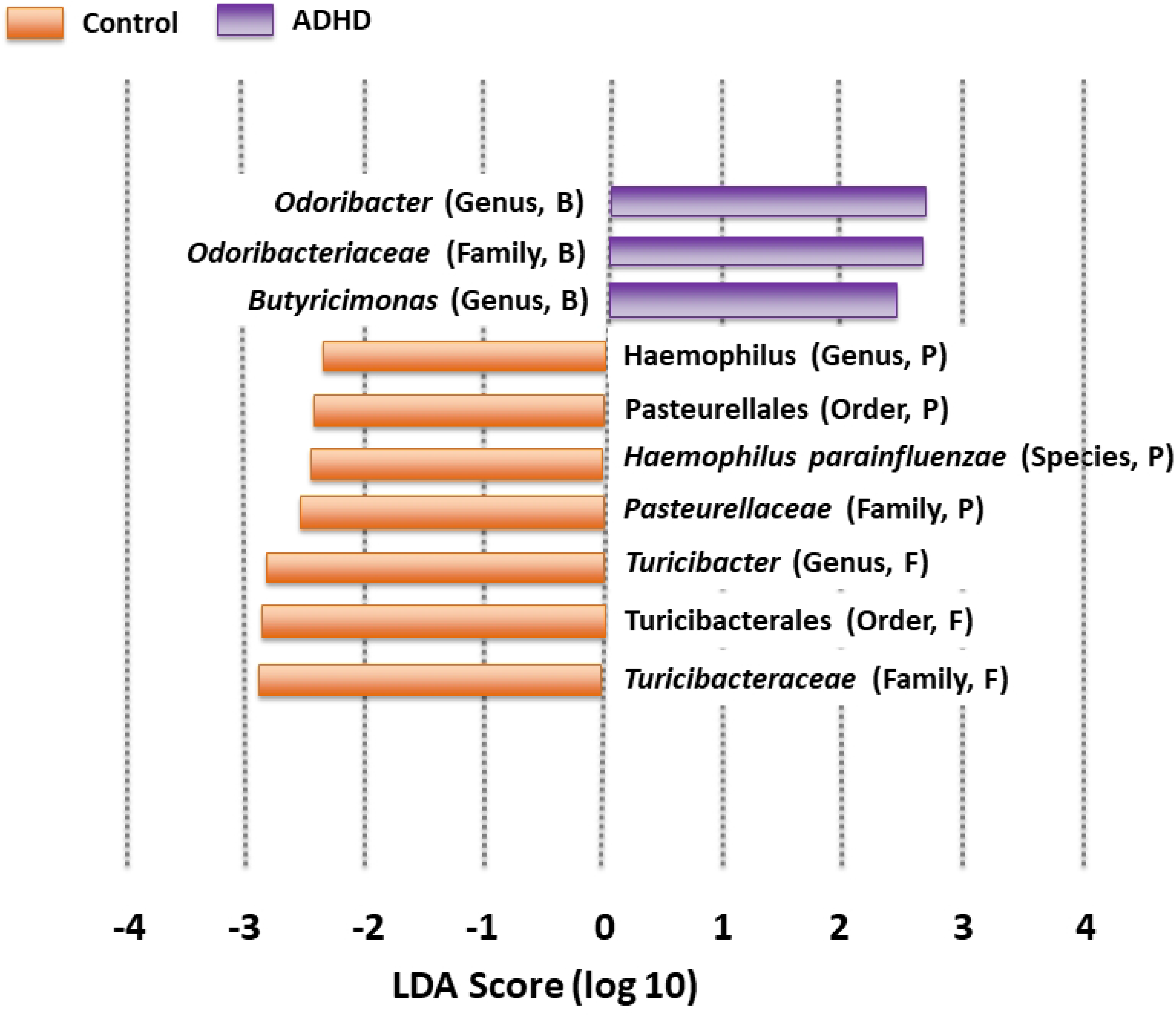
Differential Abundance. Distinguishing taxa for Control (orange) and ADHD (purple) groups, produced by LEfSe (69). Corresponding phyla for each taxon are indicated in parentheses, with B=Bacteroidetes, F=Firmicutes, and P=Proteobacteria (no distinguishing Actinobacteria were found). (a) Distinguishing taxa plotted on a cladogram, with each concentric circle representing a phylogenetic classification level (innermost=phylum). Shared areas represent distinctive regions of the phylogenetic tree. (b) Distinguishing taxa ordered by Linear Discriminant Analysis (LDA, (92)). A higher magnitude indicates more reliable differentiation.

The cladogram (Fig. 3A) shows proposed biomarkers on the phylogenetic tree, highlighting those closely related. Fig. 3A shows one Bacteroidetes family (*Odoribacteriaceae*) distinguishing ADHD, while Firmicutes (*Turicibacteriaceae* and its order Turicibacterales) and Proteobacteria (*Pasteurellaceae* and its order Pasteurellales) distinguished Control. The bar graph (Fig. 3B) uses Linear Discriminant Analysis (LDA, (92)) to order by differentiation degree, expanding to include genera and species. ADHD continues to be predominated by Bacteroidetes and includes the only two *Odoribacteriaceae* genera in our samples, *Odoribacter* and *Butyricimonas*, supporting earlier claims of *Odoribacteriaceae* as an ADHD biomarker (59). Control continues to be predominated by Firmicutes (now including *Turicibacter*) and Proteobacteria (now including *Haemophilus* and *H. parainfluenzae*).

*Haemophilus* was found as a Control biomarker by another study (57). *H. parainfluenzae,* the only *Haemophilus* species present, is a well-known lung pathogen (93), though its gut functionality remains largely unknown. Its elevated Control abundance relative to ADHD is indeed mysterious, though upon further inspection is still very low (< 0.1%).

*Turicibacter*, although never previously reported as a biomarker in an ADHD gut microbiome study, has been reported in one involving depression in mice (33). Metabolically in mice, *Turicibacter* signals the gut to produce serotonin (5-HT) (94), which influences ER (95). Both ADHD and depression are characterized by ER neurocircuitry deficiencies. LEfSe did not report any EF-associated biomarkers. This may be largely because EF is more strongly regulated by dopamine (95), for which the gut only produces roughly 50% (96), compared to 90% of 5-HT (20).

#### Compositional

Compositional analyses compare taxa relative abundances (97). We generated compositional bar charts at all phylogenetic tree levels beginning with phylum (Fig. 4A). Samples on the *x*-axis are ordered by increasing ASRS score, and the *y*-axis represents relative abundance.

**Fig. 4.**
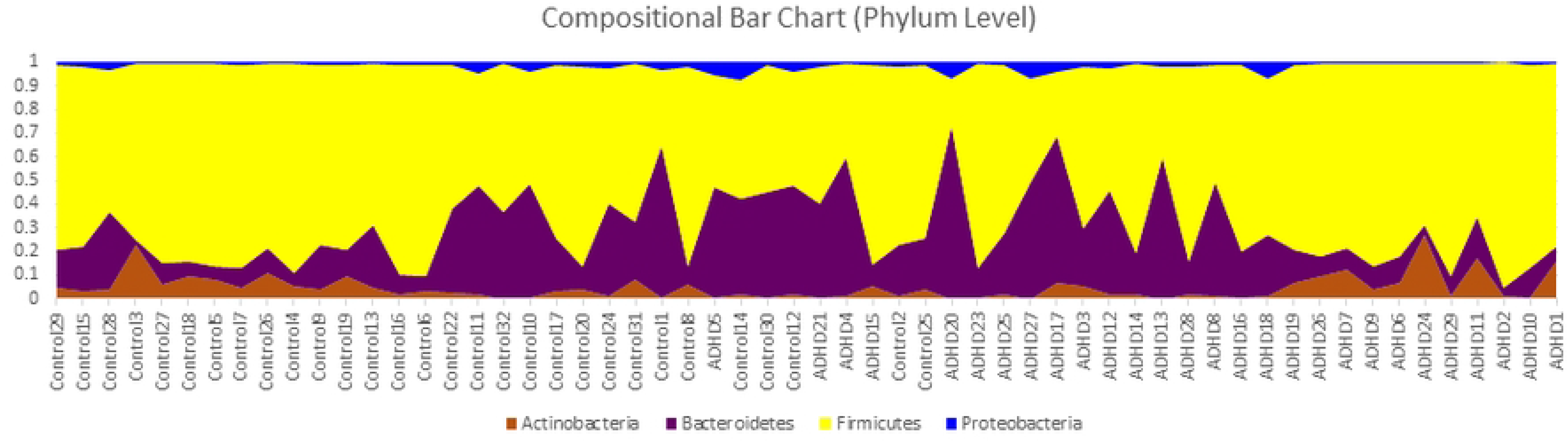
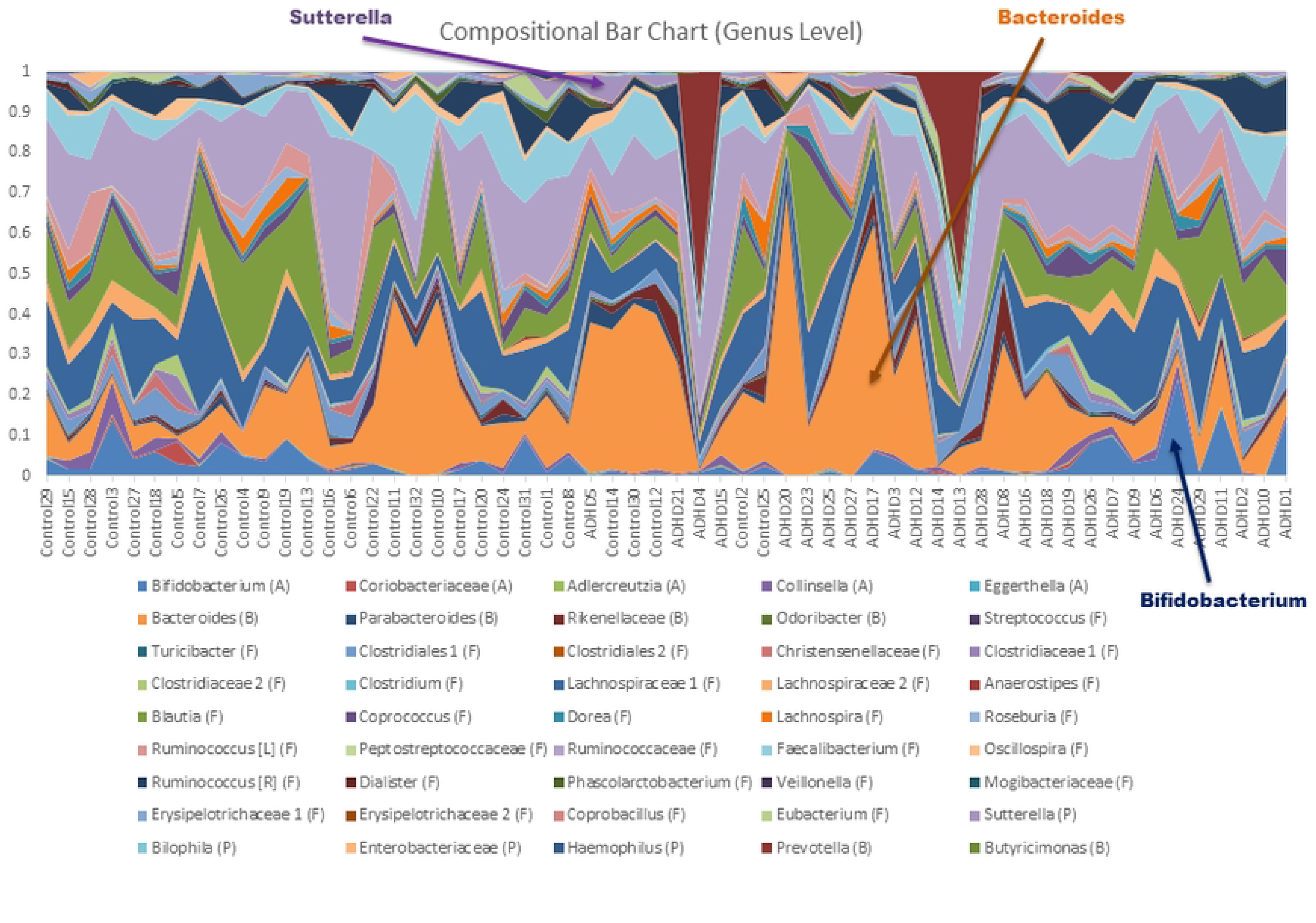
Compositional Analysis, Phylum and Genus Levels. Microbial compositional bar graph for each subject, generated using QIIME (70), conducted at (a) the phylum level and (b) the genus level. Subjects are ordered by increasing Adult ADHD Self Report Scale (ASRS) score, with the y-axis representing relative abundance.

A typical gut microbiome profile (98) is observed, dominated by Firmicutes and Bacteroidetes, followed by Actinobacteria and Proteobacteria. Control has slightly elevated Firmicutes (70-66%), mirroring an earlier study (62) that importantly (99) also sequenced the same 16S V3-V4 region. Slightly contrary to this same study, which reported this difference to be largely occupied by an ADHD Actinobacteria increase, ours was mostly occupied by an ADHD Bacteroidetes increase (from 22% to 25%). Yet Actinobacteria remains mysterious in Fig. 4A, elevated at very high ASRS scores, but also at very low scores. Bacteroidetes and Proteobacteria also appear reduced at these same extremes. These seemingly contradictory results create challenges in drawing meaningful conclusions with respect to role(s) played by these phyla. Yet they capture our interest, especially given the earlier reported anomalous behavior of an Actinobacteria genus, *Bifidobacterium*, at high and low ASRS-IV scores (58).

Class and order levels produced bar charts similar to Fig. 4A; we include these as Supplemental Fig. S2 and S3. Levels below order often had too many taxa to clearly view dynamics. We include the genus level (Fig. 4B), family and species as supplemental Fig. S4 and S5), as the genus level includes *Bifidobacterium*. And indeed, it turns out, *Bifidobacterium* (blue, bottom) has elevated abundances high and low ASRS scores, appearing most responsible for this same behavior in its phylum Actinobacteria (Fig. 4A). *Bacteroides* (orange, middle) is a highly abundant taxon that also mirrors the behavior of its phylum (Bacteroidetes, Fig. 4A), increasing in the middle and decreasing at extremes. Proteobacteria is more difficult to observe given its low relative abundance (1-2%), though *Sutterella* (lilac, top) also appears to follow this trend. All three observations are verified in supplementary Fig. S6.

This is not the first time these taxa have generated interest. Many Actinobacteria, and especially *Bifidobacterium*, have been used as probiotics and are considered elements of a health gut (100–105). As discussed, *Bacteroides* and its family *Bacteroidaceae*, as well as several member species, have been reported differentially abundant in ADHD (53,54,59,60); some elevated, others reduced. Some have argued *Bacteroides* to be the most important “window” to understanding the human gut (106). *Sutterella stercoricanis* was also reported as an ADHD biomarker (54). These same taxa make multiple appearances in studies involving other NPDs as well. *Bacteoridaceae* was the top LEfSe Major Depressive Disorder (MDD) biomarker in one study (36). Another reported elevated *Bacteroides* and reduced *Bifidobacterium* in anxiety (107). *Sutterella* is elevated in Autism Spectrum Disorder (ASD) (108), a condition with so much symptomatic overlap with ADHD that an ASD+ADHD phenotype has been established (109).

## Discussion

These analyses produced a few interesting preliminary observations, but their birds-eye view limited the depth we could pursue. Compositional analysis was a perfect example: even though there was a visible trend between ASRS score and *Bifidobacterium*, *Bacteroides,* and *Sutterella* abundances, no definitive conclusions could be produced. Fundamentally macroscale behaviors emerge from microscale interactions. We attempt to unlock some of these mysteries by now exploring *ecological relationships*.

Microbial ecological relationships take many forms. They can be positive or negative, mutual (cooperation (65,66) or competition (67)) or one-way (commensalism (110) or amensalism (111)). In particular, two-way relationships (cooperation and competition) can be approximated using correlations (73). We use SparCC (83) compute correlations, which has advantages in reducing compositional effects within relative abundances. We also use a *p*-value threshold of 0.01 to only count correlations with the highest confidence, as the historically accepted threshold of 0.05 has come under recent question (112,113). We build Microbial Co-occurrence Networks (MCNs, (72)) using taxa as nodes and SparCC correlations as edges, and perform community detection on these networks using the clustering algorithm Affinity Propagation (AP, (74)). Finally, we use Ablatio Triadium (ATria, (75)) as a centrality algorithm to produce a ranked list of important taxa in each MCN. ATria is specifically designed for signed and weighted networks, incorporating both social network (88) and economic theory (87) in its calculations. It is also iterative, removing dependencies of a central node before computing the next most central.

During our analyses we sometimes use “cooperation” to refer to a positive SparCC correlation and “competition” when referring to a negative. We emphasize, however, that correlations are an *estimate* of ecological relationships, that ultimately require further downstream analysis (multi-omics and experimental verification) before establishing official conclusions. With the underlying web of interactions being exponentially complex and large-scale laboratory experiments potentially costly, our results can provide guidance regarding target taxa and avenues to pursue.

### Part II. Ecological Relationships

#### Upper Levels: Phylum, Class, and Order

Fig. 5 shows MCNs at the phylum (Fig. 5A-B), class (Fig. 5C-D), and order (Fig. 5E-F) levels. Taxa (nodes) in all MCNs are colored by phylum (legend at the bottom of Fig. 5). Node size is proportional to relative abundance (larger=higher). Correlation (edge) color represents sign; green indicates positive (est. cooperation) and red indicates negative (est. competition). Edge thickness is proportional to correlation magnitude (thicker=stronger). Networks are visualized using the Fruchterman-Reingold algorithm (85), which spatially orients nodes based on edge weight (closer=more positive). Nodes are labeled with their taxon and provided with ATria centrality ranking if found important (format: *#rank*, T=Tie). At the phylum level only (Fig. 5A-B), we label each edge with its correlation value. Phylum-level MCNs (Fig. 5A-B) show SparCC appears to handle compositional effects well, as despite collectively encompassing about 95% of both populations, Firmicutes and Bacteroidetes are only weakly negatively correlated.

**Fig. 5.**
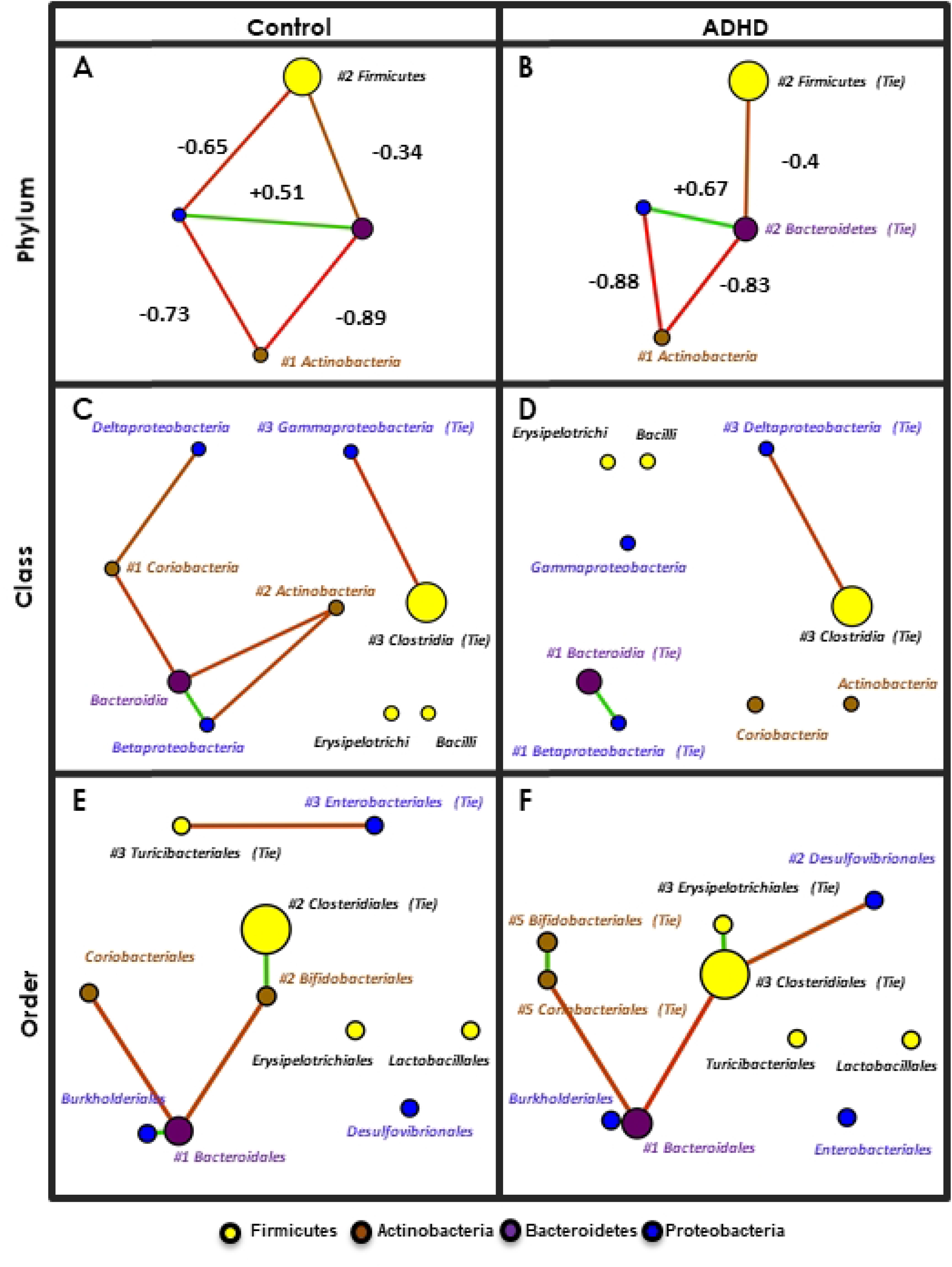
Upper-Level Microbial Co-occurrence Networks (MCNs). MCNs at the phylum (A), class (B), and order (C) taxonomic levels, visualized using Cytoscape [57], and oriented by Fruchterman-Reingold [58]. Nodes represent taxa, colored by phylum with size directionally proportional to abundance. The co-occurrences are distinguished by those that co-habit (green edges) and co-avoid (red edges). SparCC (83) correlation (p=0.01) was used as edge weight and also the parameter for Fruchterman-Reingold when determining edge length (larger=closer). SparCC correlations are shown at the phylum level. All taxa found as important by ATria are denoted by a pound sign (#) followed by its rank (ties indicated).

Table 2 shows every correlation in all three MCNs, and its sign, + (green) or – (red). Correlations that appear only in Control are highlighted orange, only in ADHD highlighted purple, and in both highlighted grey. Correlations at each taxonomic level are grouped by their next highest level classification; for example in row 1: phyla Actinobacteria and Bacteroidetes were negatively correlated in both phylum-level MCNs (Fig. 5A-B), member classes Actinobacteria and Bacteroidia were negatively correlated only in Control (Fig. 5C), as were member orders Bifidobacteriales and Bacteroidales (Fig. 5E). White, italicized correlations were not present in either MCN, but a correlation among descendants was; for example in row 3: phyla Actinobacteria and Firmicutes were not correlated in either MCN, nor were member classes Actinobacteria and Clostridia, but member orders Bifidobacteriales and Clostridiales were positively correlated in Control (Fig. 5E).

**Table 2.**
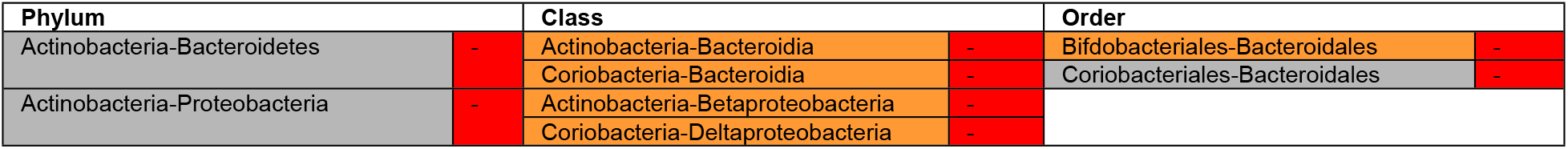
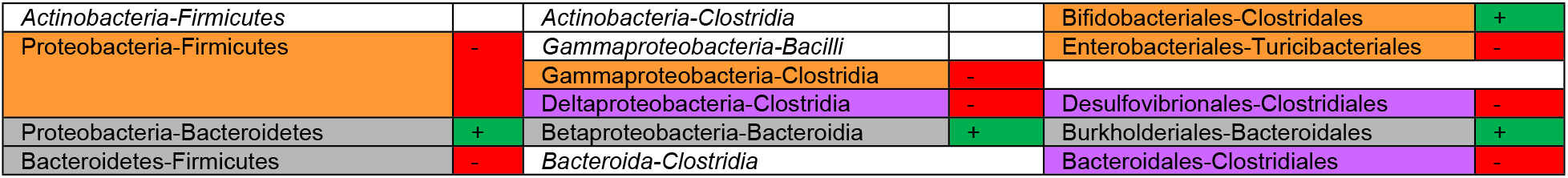
Upper-level taxa correlations, grouped by taxonomic classification.

Table 3 shows collective ATria results, similarly grouped. At each level, taxa found equally important in both MCNs are highlighted grey; taxa found more important in Control light orange, and only important in Control dark orange (analogous case for ADHD and purple). Taxa ranked as first or tied for first in either MCN are **bold**.

**Table 3.** Upper-level ATria results, grouped by taxonomic classification.

| Phylum | Class | Order |
| --- | --- | --- |
| Actinobacteria (#1/#1) | Actinobacteria (#2/NR) | Bifidobacteriales (#T2/T5) |
|  | **Coriobacteria (#1/NR)** | Coriobacteriales (NR/#T5) |
| Bacteroidetes (NR/#T2) | **Bacteroidia (NR/#T1)** | **Bacteroidales (#1/#1)** |
| Firmicutes (#2/#T2) | Bacilli (NR/NR) | Turicibacteriales (#T3/NR) |
|  | Clostridia (#T3/#T3) | Clostridiales (#T2/#T3) |
|  | Erysipelotrichia (NR/NR) | Erysipelotrichiales (NR/#T3) |
| Proteobacteria (NR/NR) | **Betaproteobacteria (NR/#T1)** |  |
|  | Deltaproteobacteria (NR/#T3) | Desulfovibrionales (NR/#2) |
|  | Gammaaproteobacteria (#T3/NR) | Enterobacteriales (#T3/NR) |

Compositional results are mirrored here: ADHD showed elevated Bacteroidetes at the expense of Firmicutes, and these taxa are negatively correlated in both MCNs (Fig. 5A-B). But while Firmicutes and Bacteroidetes dominate both populations (largest nodes, Fig. 5A-B) as is typical in the gut microbiome (98), SparCC and ATria estimate a far less abundant phylum, Actinobacteria (roughly 4% of both populations), as most important to their overall gut ecology. In both MCNs (Fig. 5A-B), phylum Actinobacteria has the strongest negative correlations and ATria ranks it first (Table 2).

We make three more observations at these upper taxonomic levels, that we keep in mind when moving to the lower:

A. **A core Proteobacteria-Bacteroidetes positive correlation (est. cooperation) forms.** Table 1 shows this, with Proteobacteria and Bacteroidetes (the only positive correlation in either phylum-level MCN), member classes Betaproteobacteria and Bacteroidia, and member orders Burkholderiales and Bacteroidiales.
B. **In Control, taxa in (A) have more negative edges with Actinobacteria (est. competition), especially Bifidobacteriales.** The highest magnitude negative edges in both phylum-level MCNs (Fig. 5A and 5B) involve Actinobacteria with Proteobacteria and Bacteroidetes. Yet while the two consistently dependent Actinobacteria classes (Actinobacteria and Coriobacteria) continue this same dynamic with Bacteroidia (Bacteroidetes) and Betaproteobacteria (Proteobacteria) in Control (Fig. 5C and Table 1), they are completely disconnected in ADHD (Fig. 5D). Worth noting, this is despite their relative abundance being nearly the same in Control/ADHD: Coriobacteria 1.5/1.1%, and Actinobacteria 3.2/3.7%. Further, ATria ranks Actinobacteria and Coriobacteria as the top two Control taxa (Table 2). In ADHD, Bacteroidia and Betaproteobacteria are the top two (Table 2), and the MCN shows no negative edges (est. competition) at all involving these taxa (Fig. 5D). The order level reveals Bifidobacteriales (Actinobacteria) may be more responsible for this difference than Coriobacteriales (Coriobacteria). While Bifidobacteriales and Coriobacteriales both continue their negative correlations with Bacteroidales (Bacteroidia) in Control, only Coriobacteriales does in ADHD. Table 1 actually shows all edges involving Bifidobacteriales to be exclusive to Control, now including a positive correlation with Clostridia (the most abundant Firmicute). An increased participation of order Bifidobacteriales thus emerges as a distinguishing feature of Control, which is further supported by ATria (Table 2), which ranks Bifidobacteriales higher (tied for second) in Control, and Coriobacteriales only in ADHD.
C. **A shift in Firmicutes-Proteobacteria dynamics.** This begins immediately at the phylum level (Fig. 5A) with Control having a negative correlation (−0.65) that is absent in ADHD (Fig. 5B). The most abundant Firmicute class (Clostridia) isnegatively correlated with different Proteobacteria classes; Gammaproteobacteria in Control (Fig. 5C), Deltaproteobacteria in ADHD (Fig. 5D), and the latter continues at the order level (Fig. 5F) with Clostridiales (Clostridia) and Desulfovibrionales (Deltaproteobacteria). In Control (Fig. 5E), a negative correlation emerges between Enterobacteriales (Gammaproteobacteria) and LEfSe Control biomarker Turicibacteriales (Bacilli).

#### Summary

Upper-level analysis revealed increased Actinobacteria participation in Control gut ecology, especially order Bifidobacteriales. Much of this involved negative correlations with a core of positively correlated Bacteroidetes (Bacteroidales) and Proteobacteria (Burkholderiales). Recalling our compositional analyses and anomalous behavior involving *Bifidobacterium* (Bifidobacteriales), *Bacteroides* (Bacteroidales), and *Sutterella* (Burkholderiales), we are now interested in exploring these dynamics at lower taxonomic levels. We will continue to observe Firmicutes-Proteobacteria dynamics, as despite a still unclear picture, a clear distinction is shown between Control and ADHD.

#### Lower Levels: Family, Genus, and Lowest Possible

Fig. 6 shows Control and ADHD MCNs at family (Fig. 6A-B), genus (Fig. 6C-D), and lowest possible taxonomic classification levels (Fig. 6E-F). In this latter MCN each taxon is classified at the species level if possible (rare with 16S), otherwise more commonly the genus level is used. Schemes regarding color, node size, and edge thickness are the same as Fig. 5. Since the MCNs are now larger we do not label every node, only those that we reference in our analyses. We also extend Table 1 to include correlations from every taxonomic level, but as this is also very large we include it as Supplemental Table S1 and extract only relevant portions to our discussion. We perform a similar task with ATria, and Supplemental Table S2.

**Fig. 6.**
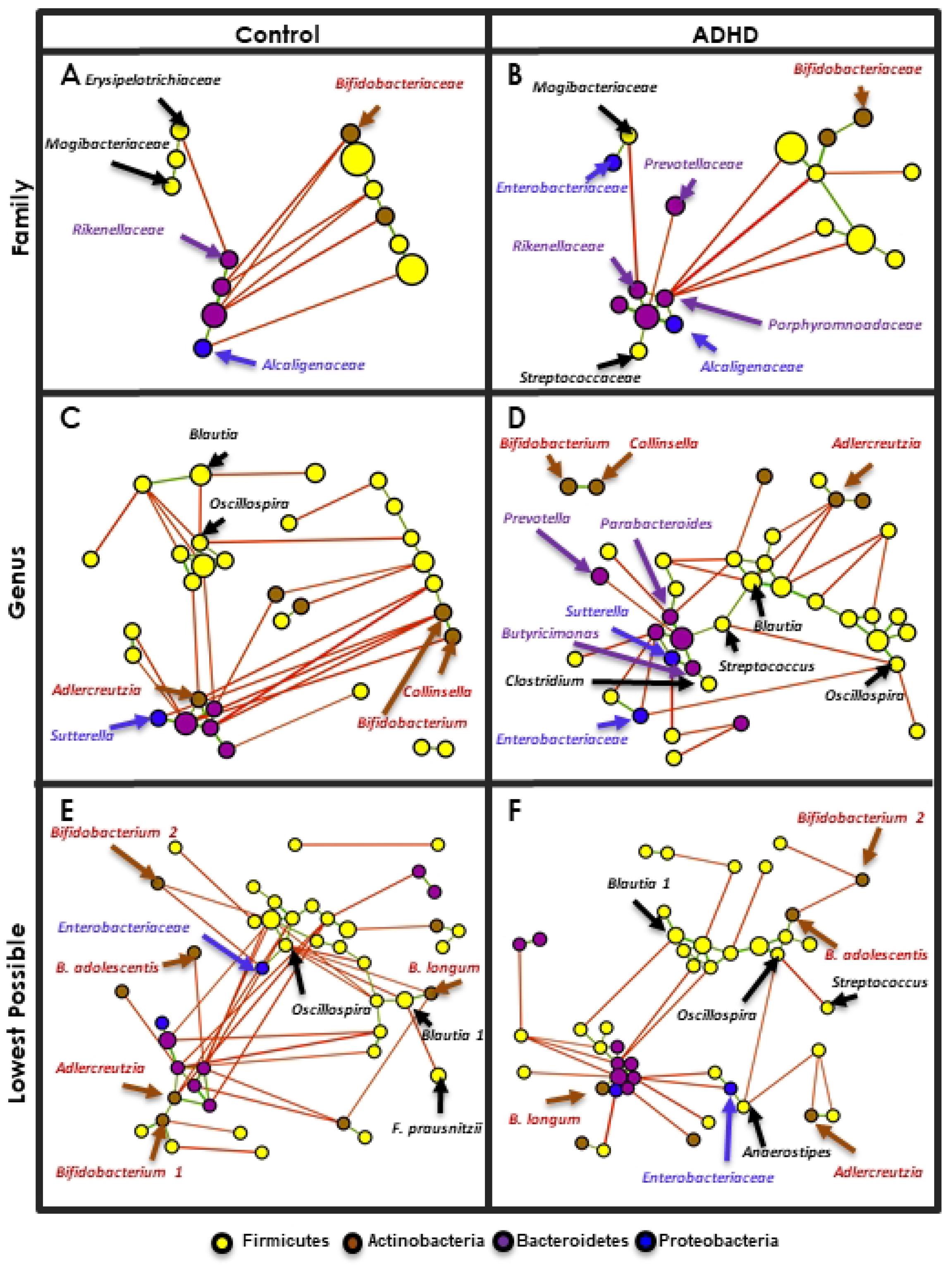
Lower-Level MCNs. MCNs at the family (A), genus (B), and species (C) taxonomic levels. Network visual properties, including node and edge size, color, and orientation, are the same as Fig. 5. Taxa noted throughout our analyses are labeled.

Fig. 6 shows taxa separating into a group of primarily Bacteroidetes (dark purple, lower left), and another of primarily Firmicutes (yellow, upper right). Enough taxa are also now present to perform meaningful community analysis. Fig. 7 shows the same MCNs as Fig. 6, after running Affinity Propagation (AP, (83)) and coloring by cluster. At the family level (Fig. 7A-B) four clusters form. One is dominated by Bacteroidetes, family *Bacteroidaceae* (*BB*, magenta). Two are dominated by Firmicutes, one family *Lachnospiraceae* (*FL*, gold), and the other family *Ruminococcaceae* (*FR*, green). In Control (Fig. 7A) the fourth cluster consists of three mixed-family Firmicutes (*FM,* dark teal). In ADHD (Fig. 7B) two of these are absent and the Proteobacteria *Enterobacteriaceae* is present, leaving it no longer Firmicutes-dominant (*M,* grey).

**Fig. 7.**
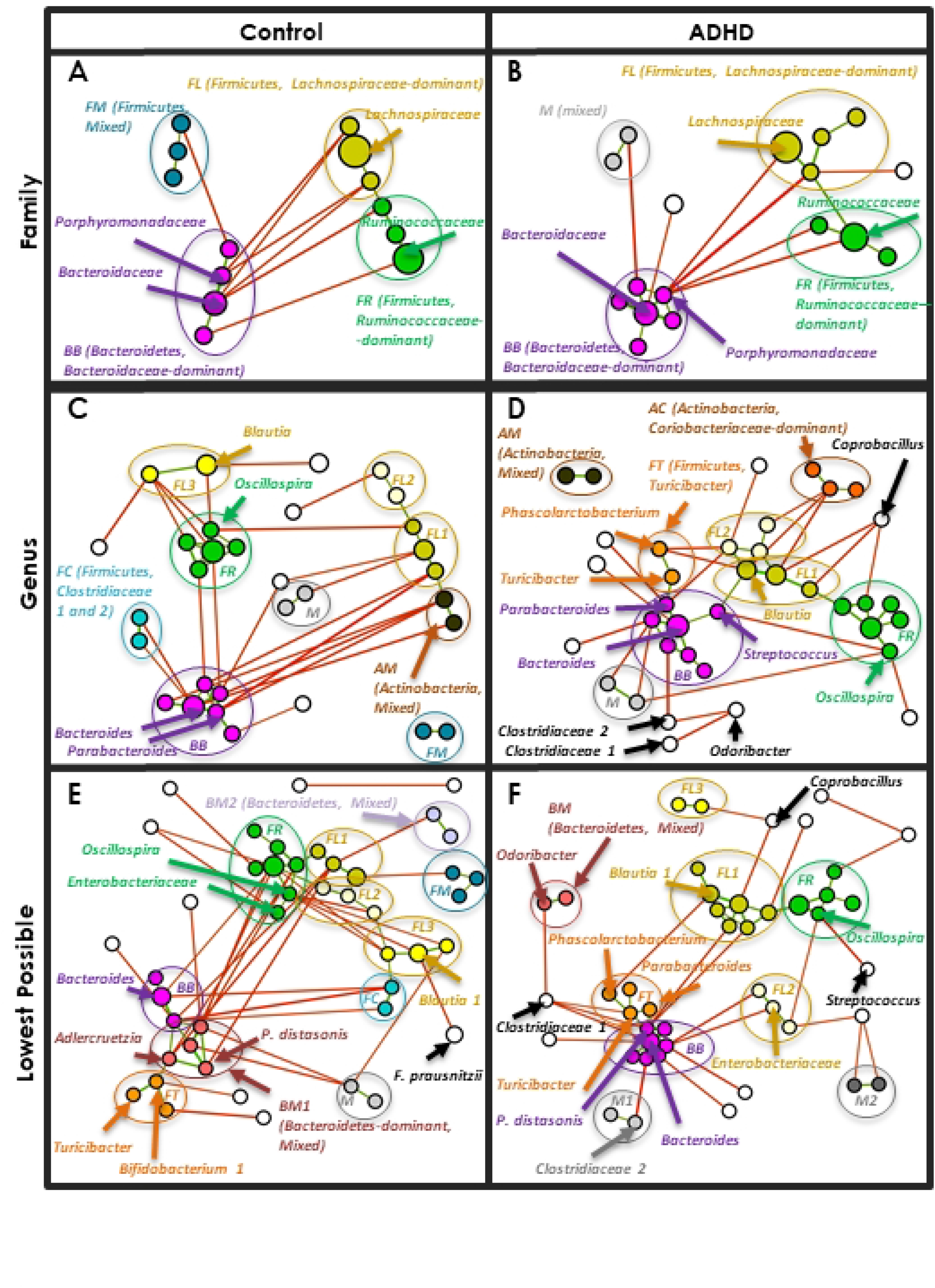
Clusters. Same MCNs as Fig. 6, after clustering with the affinity propagation (AP) algorithm (86). Family-level clusters are each given a unique color, and labeled with their dominant phylum and member family. New clusters that form at each lower taxonomic level are labeled, colored with shades corresponding to their dominant phylum/family when applicable - i.e. at the genus level *FL1*-*FL3* are different shades of gold (family-level *FL*). Taxa noted throughout our analyses are labeled.

Clusters *BB* and *FR* remain at the genus level (Fig. 7C-D). Several Firmicutes, *Lachnospiraceae*-dominant clusters emerge, referred to as *FL1*, *FL2*, etc. (gold shades). A mixed-family Actinobacteria cluster of *Bifidobacterium* and *Collinsella* forms in both MCNs (*AM*, brown), and an Actinobacteria, *Coriobacteriaceae*-dominated cluster forms in ADHD (Fig. 7D, *AC*, burnt sienna). A small group of two *Clostridiaceae* composes cluster *FC* in Control (Fig. 7C, aqua). In ADHD (Fig. 7D), a cluster (orange) emerges as the only Firmicutes-dominant cluster with positive correlations to cluster *BB*. This eventually becomes present in both lowest-level MCNs (Fig. 7E-F) with core member Control LEfSe biomarker *Turicibacter*, so we call this cluster *FT*.

At the lowest level we kept cluster names as consistent as possible with genus-level membership (for example, a cluster mostly comprised of *FL2* genus-level taxa would also be named *FL2* at the lowest level). Both MCNs (Fig. 7E-F) now include a mixed-family, Bacteroidetes-dominant cluster *BM1* (pink), and Control includes a second (*BM2,* orchid). Supplemental Tables S3-S5 list all clusters, members, and centroids at all levels. As with earlier tables, we will extract portions relevant to our discussion.

Finally, to measure cluster size, tightness, and interactions, we produce a heatmap of taxa correlations (Fig. 8) with taxa ordered on the *x*- and *y*-axes by Fig. 7 cluster. Green/red intensity at each point (*x*, *y*) denotes the degree of positive/negative correlation between taxa *x* and *y* (symmetric, by definition). Clusters appear as rough squares of positive (green) correlations on the diagonal. We outline each box with the same color as its corresponding Fig. 7 cluster.

**Fig. 8.**
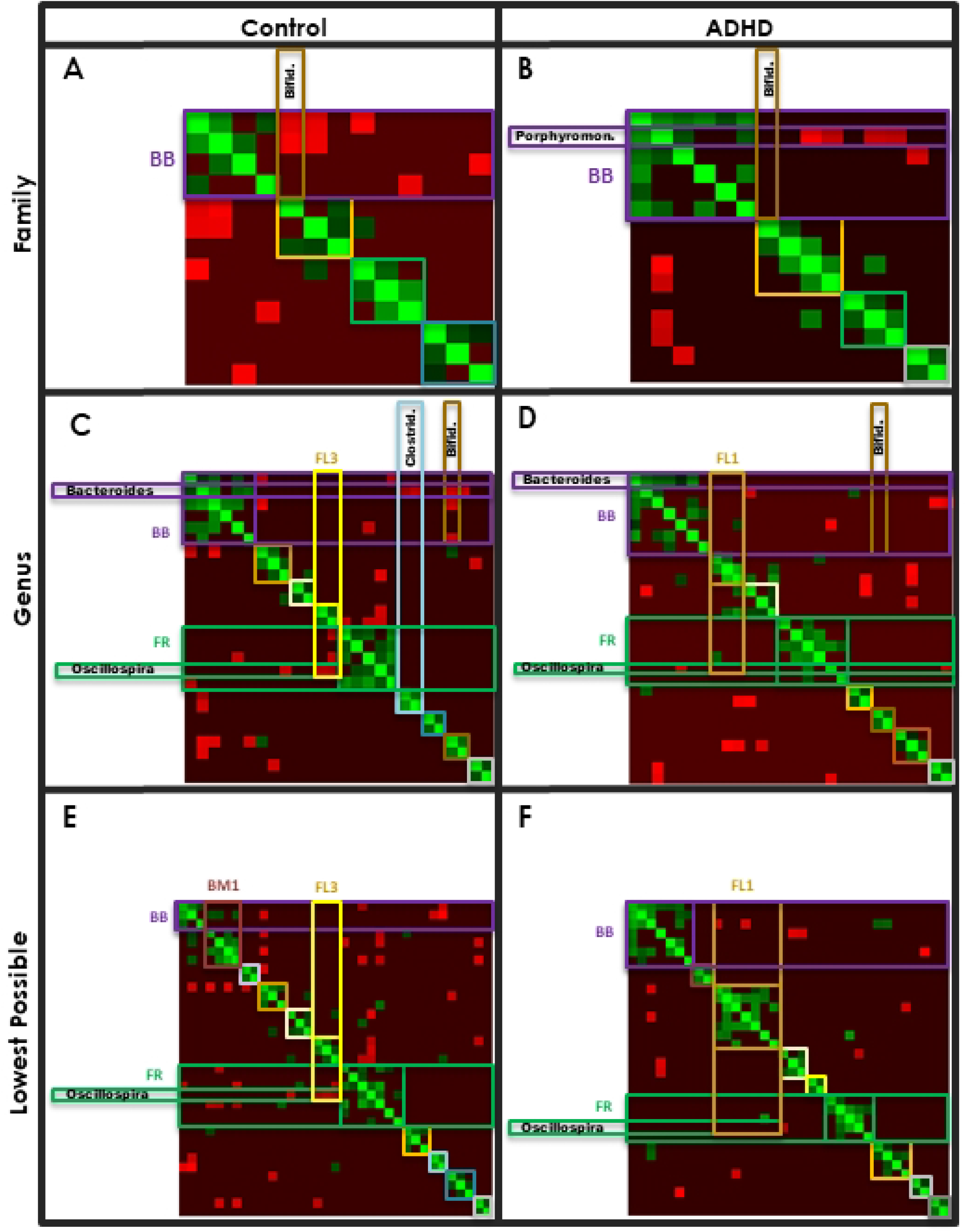
Heatmaps. Heatmap representation of taxa correlations (green=positive, red=negative), with taxa organized on each axis by cluster (symmetric matrix). The area corresponding to the intersection of each cluster with itself is outlined with a box using the corresponding cluster color in Fig. 7. Taxa and clusters noted throughout our analyses are labeled on the axes.

We first continue to pursue observations (A)-(C) from the upper taxonomic levels. Afterwards, we discuss any new and interesting trends.

(A) **A core Proteobacteria-Bacteroidetes positive correlation (est. cooperation) forms.** Recall the orders involved in this correlation were Burkholderiales (Proteobacteria) and Bacteroidales (Bacteroidetes). This corresponds to cluster *BB*, with genus *Sutterella* and multiple *Bacteroidales* taxa. In ADHD this cluster is larger and includes more *Bacteroidales* plus some Firmicutes, and nearly all members are positively correlated with its centroid *Bacteroides*. Additionally it has fewer negative correlations (est. **competition) with other clusters.**

Cluster *BB* is the only cluster with Burkholderiales and Bacteroidales descendants. Table 4 shows all correlations involving Burkholderiales and Bacteroidales lineages, organized and shaded using the same scheme as Table 1. One core positive correlation survives all six taxonomic levels in Control and ADHD (12 MCNs total, the only correlation in our entire dataset with this property). This occurs between genera *Sutterella* and *Bacteroides*. Several others involving *Sutterella* and its family *Alcaligenaceae* with cluster *BB* members are present only in ADHD – support for a larger cluster *BB* in ADHD. *Alcaligenaceae/Sutterella* are immediately visible in Fig. 6, as the only Proteobacteria (royal blue) among a slew of Bacteroidetes (dark purple).

**Table 4.**
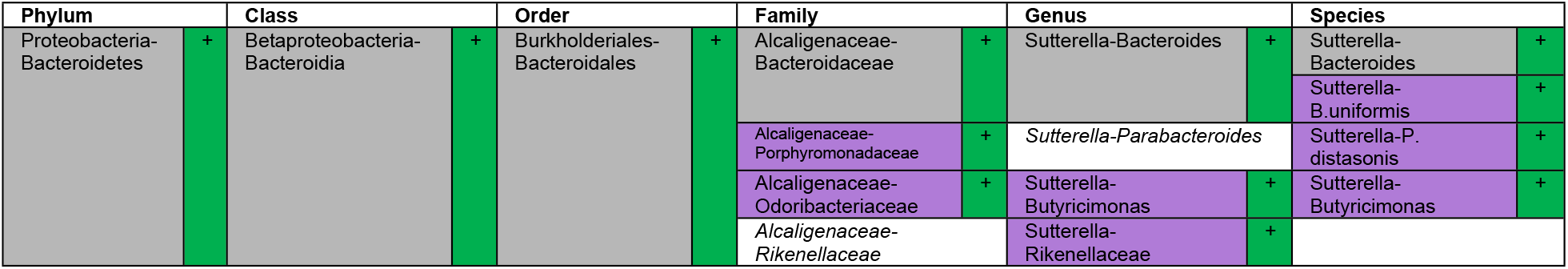
Correlations between Burkholderiales-Bacteroidales lineages, shaded using the same scheme as Table 1 (grey present in both MCNs, purple only ADHD).

Fig. 7 also illustrates the increase in ADHD cluster *BB* size, as do the heatmaps (Fig. 8, magenta square). Table 5 quantifies differences in node and edge count.

Table 5 shows cluster *BB* size to mysteriously drop in Control from the genus to the lowest level, from six taxa down to three. A closer look at Fig. 7C and 7E shows several genus-level *BB* members may be joining a mixed-family, Bacteroidetes-dominant cluster (*BM1*, pink) at the lowest level. Table 6, which shows *BB* and Control *BM1* members, confirms this. Core *BB* members are shown in bold, while italicized members are unique to Control or ADHD. Taxa of genus-level Control cluster *BB* members *Odoribacter*, *Adlercruetzia*, *Parabacteroides* (*P. distasonis*) and *Bacteroides* (*B. ovatus*) compose Control cluster *BM1* at the lowest level.

**Table 5.**
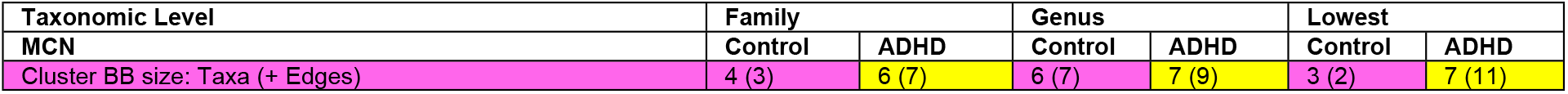
Control and ADHD cluster *BB* size. Notation: *Taxa (edges)*.

| Taxonomic Level | Family |  | Genus |  | Lowest |  |
| --- | --- | --- | --- | --- | --- | --- |
| MCN | Control | ADHD | Control | ADHD | Control | ADHD |
| Cluster BB size: Taxa (+ Edges) | 4 (3) | 6 (7) | 6 (7) | 7 (9) | 3 (2) | 7 (11) |

**Table 6.** Bacteroides, *Bacteroidaceae* dominant clusters (*BB*) and Bacteroidetes, Mixed family (*BM1*) cluster in Control. Core taxa are bold, taxa exclusive to one MCN (Control or ADHD) are italicized, and centroids are marked with an asterisk (*).

| Level | Community | Cluster Type | Control Taxon | Phylum | ADHD Taxon | Phylum |
| --- | --- | --- | --- | --- | --- | --- |
| Family | Bacteroidetes-dominant (B) | Bacteroidaceae-dominant (BB) | Bacteroidaceae |  | Bacteroidaceae* |  |
|  |  |  | Porphyromonadaceae* |  | Porphyromonadaceae |  |
|  |  |  | Alcaligenaceae |  | Alcaligenaceae |  |
|  |  |  | Rikenellaceae |  | Rikenellaceae |  |
|  |  |  |  |  | Odoribacteraceae |  |
|  |  |  |  |  | Streptococcaceae |  |
| Level | Community | Cluster Type | Taxon | Phylum | Taxon | Phylum |
| Genus | Bacteroidetes-dominant (B) | Bacteroidaceae-dominant (BB) | Bacteroides |  | Bacteroides* |  |
|  |  |  | Parabacteroides* |  | Parabacteroides |  |
|  |  |  | Sutterella |  | Sutterella |  |
|  |  |  | Rikenellaceae |  | Rikenellaceae |  |
|  |  |  | Odoribacter |  | Butyricimonas |  |
|  |  |  | Adlercruetzia |  | Streptococcus |  |
|  |  |  |  |  | Clostridium |  |
| Level | Community | Cluster Type | Taxon | Phylum | Taxon | Phylum |
| Species | Bacteroidetes-dominant (B) | Bacteroidaceae-dominant (BB) | Bacteroides* |  | Bacteroides* |  |
|  |  |  | Bacteroides uniformis |  | Bacteroides uniformis |  |
|  |  |  | Sutterella |  | Sutterella |  |
|  |  |  |  |  | Parabacteroides distasonis |  |
|  |  |  |  |  | Rikenellaceae |  |
|  |  |  |  |  | Butyricimonas |  |
|  |  |  |  |  | Bifidobacterium longum |  |
|  |  | Bacteroidetes, Mixed (BM1) | Odoribacter |  |  |  |
|  |  |  | Bacteroides ovatus |  |  |  |
|  |  |  | Parabacteroides distasonis* |  |  |  |
|  |  |  | Adlercruetzia |  |  |  |

Table 7 supports weakened connections between *BB* and *BM1* taxa in Control, showing higher intra-correlation values (0.61 and 0.62) relative to inter-correlation (0.44).

**Table 7.** Cluster *BB* and Control *BM1* intra- and inter-correlations.

| Control BB (Intra) |  |  | Control BM1 (Intra) |  |  | Control BB-BM1 (Inter) |  |  | ADHD BB (Intra) |  |  |
| --- | --- | --- | --- | --- | --- | --- | --- | --- | --- | --- | --- |
| Taxa | Edges | Mean Correlation | Taxa | Edges | Mean Correlation | Taxa | Edges | Mean Correlation | Taxa | Edges | Mean Correlation |
| 3 | 2 | 0.61 ± 0.15 | 4 | 4 | 0.62 ± 0.1 | 4 | 3 | 0.44 ± 0.02 | 7 | 11 | 0.56 ± 0.12 |

Table 6 also shows cluster *BB* members that differ between the MCNs. Cluster *BB* gains a different Actinobacteria – *B. longum* (ADHD) and *Adlercruetzia* (Control, eventually joining *BM1*). The presence of Firmicutes (yellow) is exclusive to ADHD, including *Streptococcaceae* and member genus *Streptococcus*, plus *Clostridium*. ADHD LEfSe biomarkers *Odoribacteriaceae* and *Butyricimonas* join cluster *BB* only in ADHD, and the sole *Clostridium* connection to cluster *BB* is with *Butyricimonas* (Fig. 6D).

Table 6 also indicates *BB*/*BM1* centroids, which we see across the board for ADHD are *Bacteroides* and its family *Bacteroidaceae*. In Control this belongs to *Porphyromonadaceae* (family) and descendant *Parabacteroides* (genus), until the *BB*-*BM1* “split” where *Bacteroides* becomes centroid of *BB* and *P. distasonis* of *BM1*. Table 8 shows connectivity of each of these taxa within their corresponding cluster. Percentagewise, in ADHD *Bacteroidaceae*/*Bacteroides* is a much stronger centroid; in fact over all levels only one cluster *BB* taxon was not positively correlated (*Clostridium*, genus level). Particularly given the ADHD cluster *BB* size increase, this could imply a significant role of *Bacteroidaceae*/*Bacteroides* in stabilizing a large ADHD Bacteroidetes-dominant community (would require additional experiments to verify).

**Table 8.**
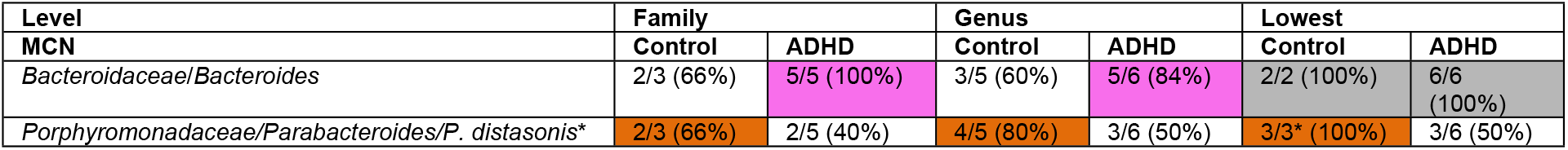
Cluster *BB* (* = *BM1*) connectivity with centroid taxa.

Interestingly ATria (Table 9) shows *Bacteroidaceae/Bacteroides* and lineages to nearly always have higher importance in Control, supporting a more “global” importance to overall gut ecology as opposed to a more local importance (cluster *BB*) in ADHD. MCNs agree, as in ADHD *Bacteroidaceae/Bacteroides* have few connections outside cluster *BB* (Fig. 7B, D, F). In Control (Fig. 7A, C, E) *Bacteroidaceae/Bacteroides* have many external connections, mostly negative (est. competition).

**Table 9.**
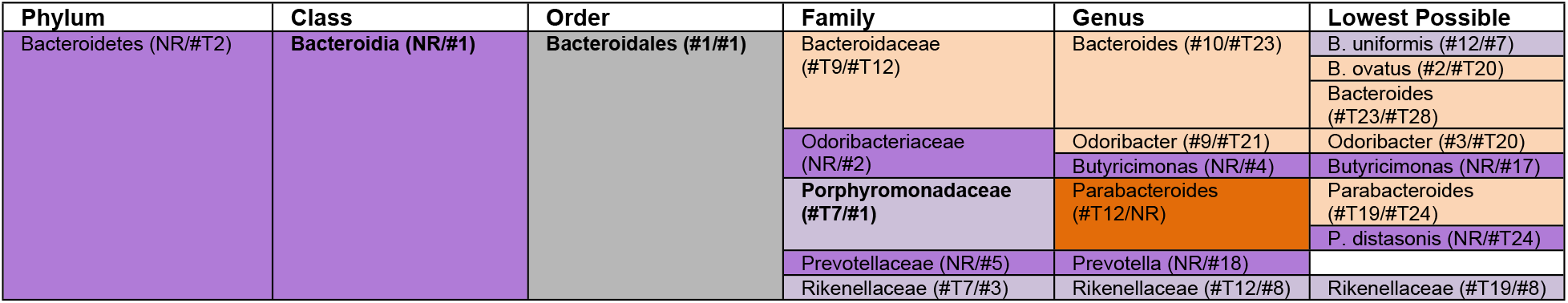
ATria rankings of Bacteroidetes taxa.

Control MCNs (Fig. 7A, C, E) and heatmaps (Fig. 8A, C, E, magenta rectangle) show negative correlations (red) to be fairly evenly distributed among cluster *BB* taxa. By contrast in ADHD (Fig. 7B, 8B), nearly all cluster *BB* negative correlations are localized to *Porphyromonadaceae* (ranked #1 by ATria). Fig. 6B shows *Porphyromonadaceae* to be the sole cluster *BB* member negatively correlated with the Firmicutes-dominant portion (Fig. 6B, upper right, collectively more than 70% of the population).

Table 10 shows that for all MCNs, in Control more than two-thirds of cluster *BB* had negative correlations with members of other clusters, compared to less than half in ADHD. Negative edge count was also almost always higher for Control, despite a smaller cluster *BB*. Collectively these results show that in Control cluster *BB* is smaller, and more connected to other clusters, primarily through negative correlations (est. competition). In ADHD cluster *BB* is larger, and more isolated.

**Table 10.**
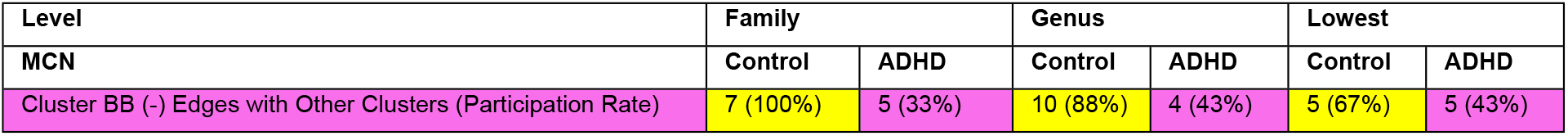
Negative correlations between cluster *BB* and other clusters. *Number (participation rate)*.

| Level | Family |  | Genus |  | Lowest |  |
| --- | --- | --- | --- | --- | --- | --- |
| MCN | Control | ADHD | Control | ADHD | Control | ADHD |
| Cluster BB (-) Edges with Other Clusters (Participation Rate) | 7 (100%) | 5 (33%) | 10 (88%) | 4 (43%) | 5 (67%) | 5 (43%) |

Table 11 provides a few final interesting observations for various Bacteroidetes taxa.

**Table 11.**
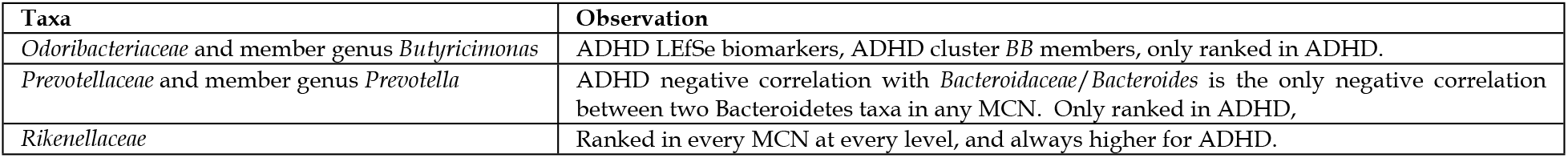
Additional observations for some Bacteroidetes taxa.

**(B) In Control, taxa in (A) have more negative edges with Actinobacteria (est. competition), especially Bifidobacteriales.** We now know taxa from (A) to correspond to cluster *BB*, which in both MCNs contained one core Proteobacteria (*Alcaligenaceae*/*Sutterella*) and otherwise primarily Bacteroidetes. We also observed cluster *BB* taxa to have far more negative correlations (est. competition) with other clusters in Control. We now see if this is also true with Bifidobacteriales lineages, including *Bifidobacterium*. Our analysis in fact reveals that **negative correlations between *Bifidobacterium* or <u>any</u> parent/descendant with <u>any</u> Bacteroidetes or Proteobacteria are exclusive to Control and absent in ADHD.**

Table 12 shows all correlations involving *Bifidobacterium* and its lineages, grouped and colored as in previous tables. Not only are negative Bacteroidetes correlations exclusive to Control (orange), but these taxa include the most abundant Bacteroidetes *Bacteroidaceae*/*Bacteroides* (ADHD cluster *BB* centroid), as well as *Porphyromonadaceae*/*Parabacteroides* (Control cluster *BB* centroid). Another appears at the lowest level between *B. adolescentis* and *B. ovatus*. With Proteobacteria, negative *Bifidobacterium* correlations are also observed with *Sutterella* (core cluster *BB* member) and *Enterobacteriaceae*, also only in Control. Heatmaps confirm *Bifidobacteriaceae*/*Bifidobacterium* to be negatively correlated with cluster *BB* taxa only in Control (Fig. 8A-D, intersection of brown and magenta rectangles). By contrast, the only ADHD correlation is positive and within cluster *BB* (*B. longum* with *B. uniformis*).

**Table 12.**
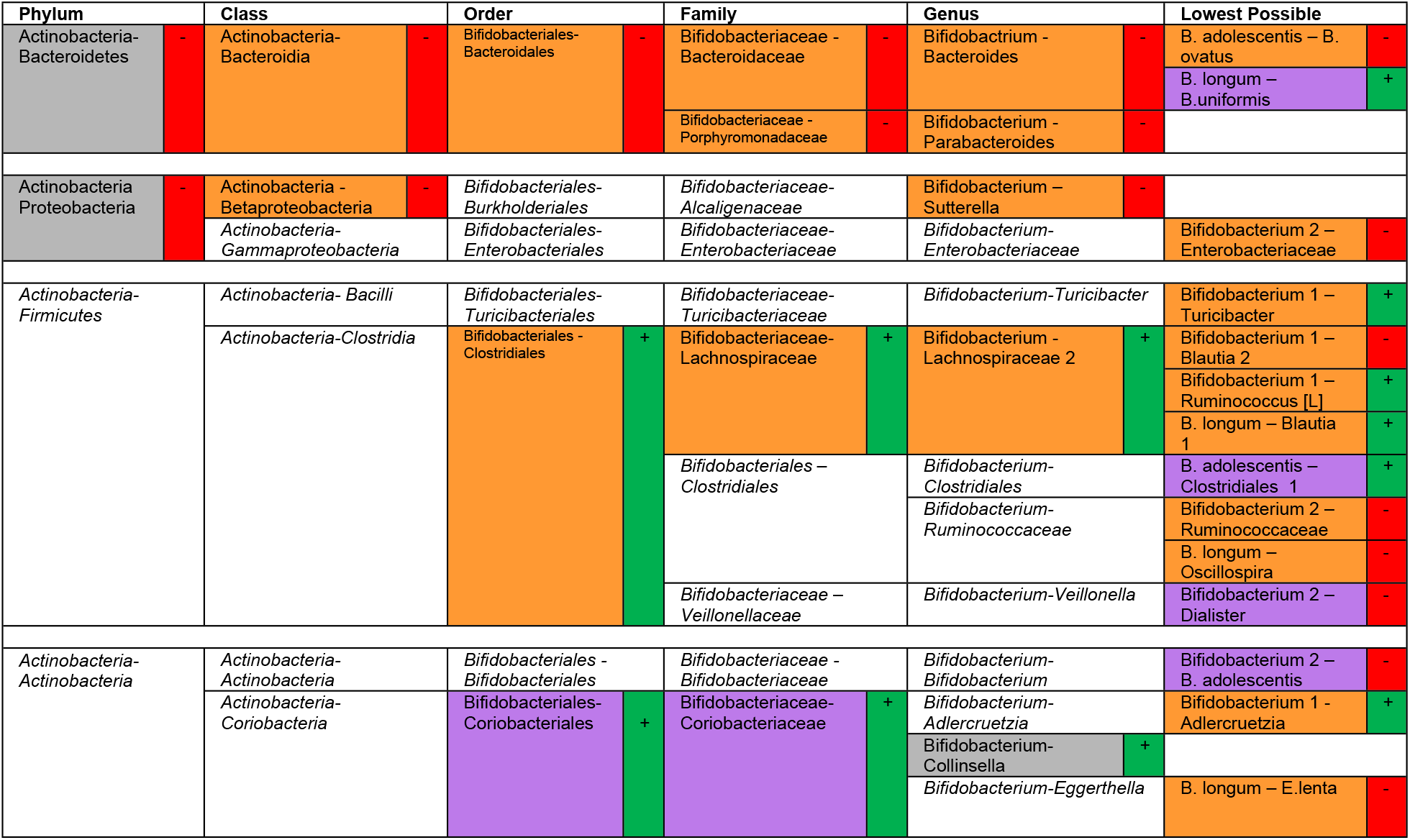
Correlations involving *Bifidobacterium* and its lineages.

Table 12 also shows *Bifidobacterium* to even have far more Firmicutes connections (positive and negative) in Control. Collectively 24 correlations were observed in Control, compared to 9 in ADHD, supporting an overall increase in *Bifidobacterium* participation in Control. ATria (Table 13) also almost uniformly ranks Bifidobacterium and its lineages higher in Control. Again, this is despite *Bifidobacterium* abundances being relatively the same (slightly higher in ADHD in fact, 3.6% to 3.2%).

**Table 13.** ATria rankings of *Bifidobacterium* and lineages.

| Phylum | Class | Order | Family | Genus | Lowest Possible |
| --- | --- | --- | --- | --- | --- |
| Actinobacteria (#1/#1) | Actinobacteria (#2/NR) | Bifidobacteriales (#T2/ #T5) | Bifidobacteriaceae (#5/NR) | Bifidobacterium (NR/#T19) | Bifidobacterium 1 (#10/NR) |
|  |  |  |  |  | Bifidobacterium 2 (#1/11) |
|  |  |  |  |  | B. longum (#15/NR) |

**(C) A Shift in Firmicutes-Proteobacteria dynamics.** Only two Proteobacteria families/genera were consistently present. One was *Sutterella* (family *Alcaligenaceae*), already noted as a core cluster *BB* member. The other is ***Enterobacteriaceae*, which our analysis supports being mostly responsible for this shift**.

Table 14 shows all Proteobacteria-Firmicutes correlations. A couple of negative correlations can be seen involving *Alcaligenaceae*/*Sutterella*, with Firmicutes *Ruminococcaceae* (Control) and *Clostridiaceae* (both). Far more significant are the differences involving *Enterobacteriaceae*. One is its negative correlation with genus *Oscillospira* in ADHD (genus level), that becomes a positive correlation with *Oscillospira* in Control (lowest level). <u>This is the only time, over all twelve MCNs, where a</u> <u>correlation sign changed between the same two taxa in Control vs. ADHD</u>.

**Table 14.**
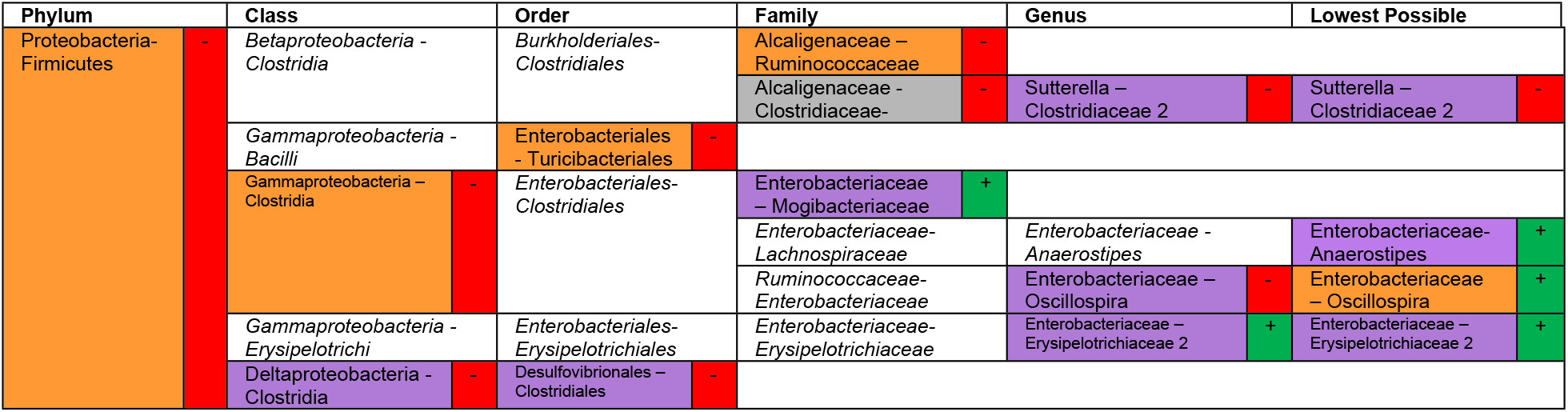
Proteobacteria-Firmicutes correlations.

Interesting shifts involving *Enterobacteriaceae* and various Firmicutes occur even at the family level, however. A small mixed-family, Firmicutes-dominant cluster *FM* forms (Fig. 7A, upper left), consisting of *Mogibacteriaceae*, *Christensenellaceae*, and *Erysipelotrichiaceae* (Table 15). In ADHD, *Enterobacteriaceae* instead joins *Mogibacteriaceae* to form a small two-taxon mixed cluster *M* (Fig. 7B, upper left, and Table 15).

**Table 15.** Mixed-family Control and ADHD clusters.

| Community | Cluster Type | Control |  |  | ADHD |  |  |
| --- | --- | --- | --- | --- | --- | --- | --- |
|  |  | Cluster | Taxon | Phy | Cluster | Taxon | Phy |
| Firmicutes-dominant (F) | Mixed (FM) | *FM* | *Mogibacteriaceae* |  |  |  |  |
|  |  | *FM* | *Christensenellaceae\** |  |  |  |  |
|  |  | *FM* | *Erysipelotrichiaceae* |  |  |  |  |
| Mixed (M) | N/A |  |  |  | *M* | *Mogibacteriaceae* |  |
|  |  |  |  |  | *M* | *Enterobacteriaceae* |  |

Dynamics of *FM* and *M* taxa change between the MCNs. Fig. 7A-B shows a distinguishing core *FM*/*M* feature is the negative correlation with *Rikenellaceae* of cluster *BB*, but the taxon involved changes from *Erysipelotrichiaceae* in Control to *Mogibacteriaceae* in ADHD. Table 16 (ATria) shows the two taxa from Control cluster *FM* “replaced” by *Enterobacteriaceae* in ADHD cluster *M*, *Christensenellaceae* and *Erysipelotrichiaceae*, are only ranked in Control, and *Mogibacteriaceae* only ranked in ADHD. This applied across all descendants, with the one notable exception being *Coprobacillus* (*Erysipelotrichiaceae*), ranked #1 for ADHD at the genus and lowest levels (the only taxon to be ranked #1 in two MCNs). We label it in Fig. 7D, F, noting its negative correlations with multiple Firmicutes-dominant clusters.

**Table 16.**
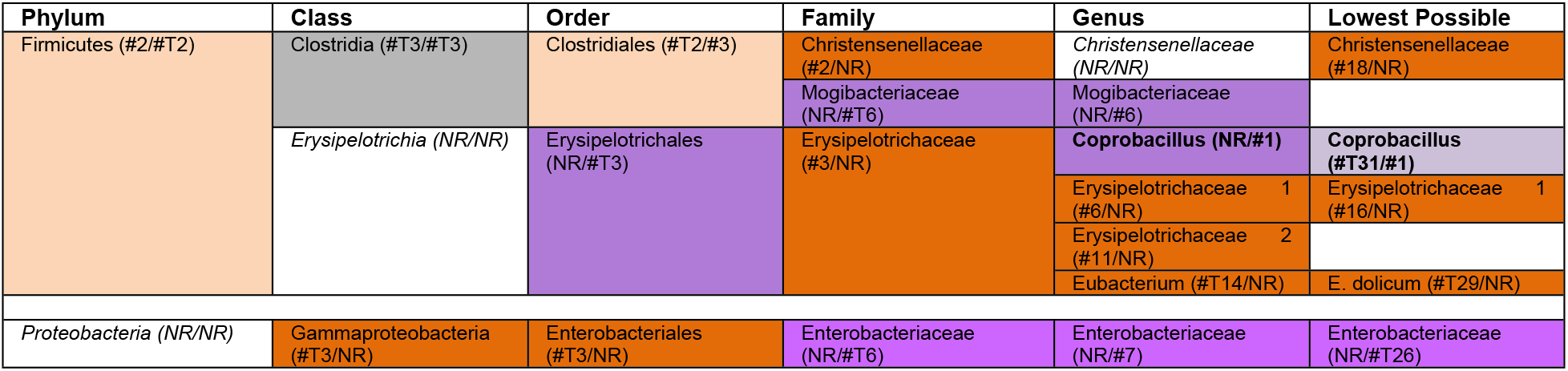
Mixed-family cluster member ATria rankings.

*Enterobacteriaceae* was also only ranked in ADHD, across all three lower levels. Its *Oscillospira* positive correlation (Table 14) is the only Control correlation involving *Enterobacteriaceae*, and *Enterobacteriaceae* actually joins *Oscillospira*’s cluster (*FR,* Fig. 7E) in Control. The sign change takes place at the genus level in ADHD (Fig. 6D), where *Oscillospira* and *Enterobacteriaceae* are negatively correlated. Although this correlation did not persist to the lowest level (Fig. 6F), *Enterobacteriaceae* is still positively correlated with *Anaerostipes*, a taxon negatively correlated with *Oscillospira* across the board. We therefore observe *Enterobacteriaceae* dynamics to shift from a state that favors *Oscillospira* <u>cooperation</u> in Control, to *Oscillospira* <u>competition</u> in ADHD. The role of *Enterobaceriaceae* in gut ecology has historically been controversial (114), with both beneficial (115) and pathogenic (116) properties emerging. Gut dysbiosis has actually been shown to trigger horizontal gene transfer between the two types (117).

#### New Observations

We make the following new observations at the lower levels.

**(D) LEfSe Biomarkers: *Turicibacter* and *Odoribacter.*** Earlier we noted ADHD LEfSe biomarkers *Odoribacteriaceae* and *Butyricimonas* as ADHD cluster *BB* members (Table 11). Biomarker *H. influenzae* and its lineages were never connected to any of our MCNs. We now observe remaining biomarkers *Turicibacter* (Control) and *Odoribacter* (ADHD).

Cluster *FT* (Fig. 7, orange) was the only Firmicutes-dominant cluster with members positively correlated with any Bacteroidetes-dominant cluster (*BB* in ADHD, *BM1* in Control). We named this cluster *FT* because of core member *Turicibacter*. *Turicibacter* (Firmicutes, LEfSe Control biomarker), which joins *Phascolarctobacterium* (Firmicutes, reduced in inattention, (57)) to form *FT* at the genus level in ADHD (Fig. 7D), where it is not present in Control. At the lowest level, *FT* is slightly larger (by one taxon) in ADHD. Supplementing the earlier trend of less cluster *BB* negative correlations (est. competition) in ADHD, this also supports the presence of a larger cluster with positive correlations (est. cooperation) as well, with *Turicibacter* as its centroid (Table 17).

**Table 17.**
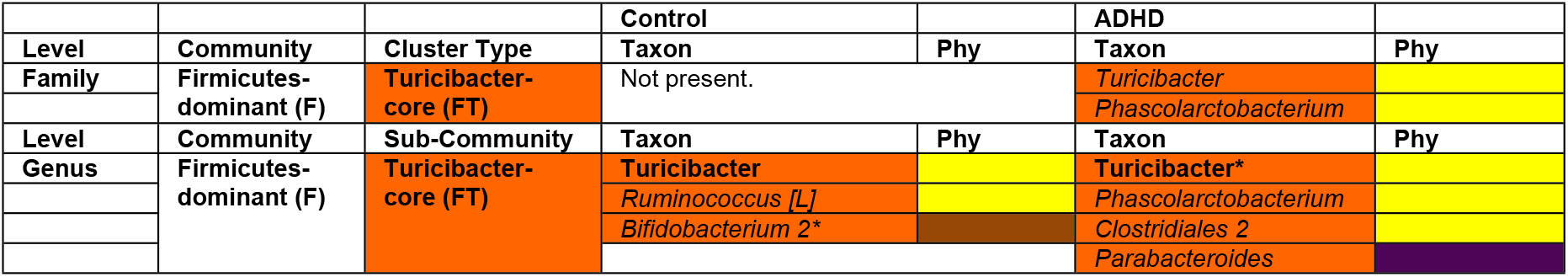
Cluster FT (Firmicutes-dominant, *Turicibacter*-core) members.

In ADHD *Turicibacter* provides the sole genus-level (Fig. 7D) *FT*-*BB* positive correlation, with *Parabacteroides* (Bacteroidetes, reported elevated in hyperactivity, (53)). At the lowest level (Fig. 7F) *Parabacteroides* joins *FT*, and along with *Turicibacter* forms *FT*-*BB* positive correlations, with member species *P. distasonis*. Interestingly in Control (Fig. 7E), the *FT*-*BB* positive correlation does not involve Firmicutes or Bacteroidetes taxa at all, but rather two Actinobacteria –*Bifidobacterium 1 (FT* centroid), and *Adlercruetzia* (*BB*). This continues our observed increases in Actinobacteria and particularly *Bifidobacterium* involvement in Control gut ecology.

Cluster *FC* forms in Control (Fig. 7C,E, aqua) and contains two *Clostridiaceae* taxa. In both MCNs these taxa negatively correlate with multiple cluster *BB* members, and in ADHD (Fig. 7F) *Clostridiaceae 1* has negative correlations with *BB* centroid *Bacteroides* plus taxa involved in *FT*-*BB* cooperation: *P. distasonis*, and *FT* centroid *Turicibacter*. In both MCNs, they participate in correlations that favor cluster *BB* competition (especially the more abundant *Clostridiaceae 1*).

Exclusive to ADHD is a negative correlation (est. competition) between these *Clostridiaceae* taxa and ADHD biomarker *Odoribacter* – both at the genus level (Fig. 7D), and *Clostridiaceae 1* at the lowest level (Fig. 7F). *Odoribacter* was reported by LEfSe as elevated in ADHD, and this negative correlation implies that an increase in *Odoribacter* abundance will decrease *Clostridiaceae 1*. Upon further inspection *Clostridiaceae 1* relative abundance is indeed reduced by a factor of two in ADHD vs. Control. *Clostridiaceae 1* and *2* cooperation in Control (forming *FC*) is also absent in ADHD.

**(E) Changes in the role of *Adlercruetzia* (Actinobacteria).** In contrast to *Bifidobacterium* (*Bifidobacteriaceae*), *Adlercruetzia* is a member of the other consistently present Actinobacteria family, *Coriobacteriaceae*. While the distinguishing feature of *Bifidobacteriaceae/Bifidobacterium* was increased Control participation, the distinguishing feature of *Coriobacteriaceae* appears to be changes in cluster membership. In fact over all *Coriobacteriaceae* descendants, only once (*Collinsella*, genus level, cluster AM, Fig. 7C-D) were any in the same Control and ADHD cluster. Table 18 also shows ATria results to be more mixed for *Coriobacteriaceae*, compared to *Bifidobacteriaceae* (Table 13).

**Table 18.** ATria rankings of *Coriobacteriaceae* and its lineages.

| Phylum | Class | Order | Family | Genus | Lowest Possible |
| --- | --- | --- | --- | --- | --- |
| Actinobacteria (#1/#1) | Coriobacteria (#1/NR) | Coriobacteriales (NR/#T5) | Coriobacteriaceae (NR/#4) | Adlercruetzia (NR/#17) | Adlercruetzia (#14/NR) |
|  |  |  |  | Collinsella (NR/#T19) | C. aerofaciens (NR/#15) |
|  |  |  |  | Coriobacteriaceae (#18/NR) |  |
|  |  |  |  | Eggerthella (#4/NR) | E. lenta (#T25/NR) |

We earlier noted *Adlercruetzia* as the Actinobacteria member of cluster *BB*/*BM1* in Control, and (along with *Bifidobacterium 1*) connecting clusters *FT* and *BB*. Table 19 shows that outside of *Bifidobacterium 1*, its positive correlations in Control were entirely with Bacteroidetes taxa (all *BB/BM1* members). By contrast in ADHD, *Adlercruetzia* relationships mostly occur with Firmicutes, including a cluster membership with *Eubacterium/E. dolicum*. Several negative correlations are seen between *Adlercruetzia* and different Firmicutes, with no overlap between Control and ADHD. This suggests *Adlercruetzia* may play a significantly different role in Control and ADHD gut ecologies.

**Table 19.**
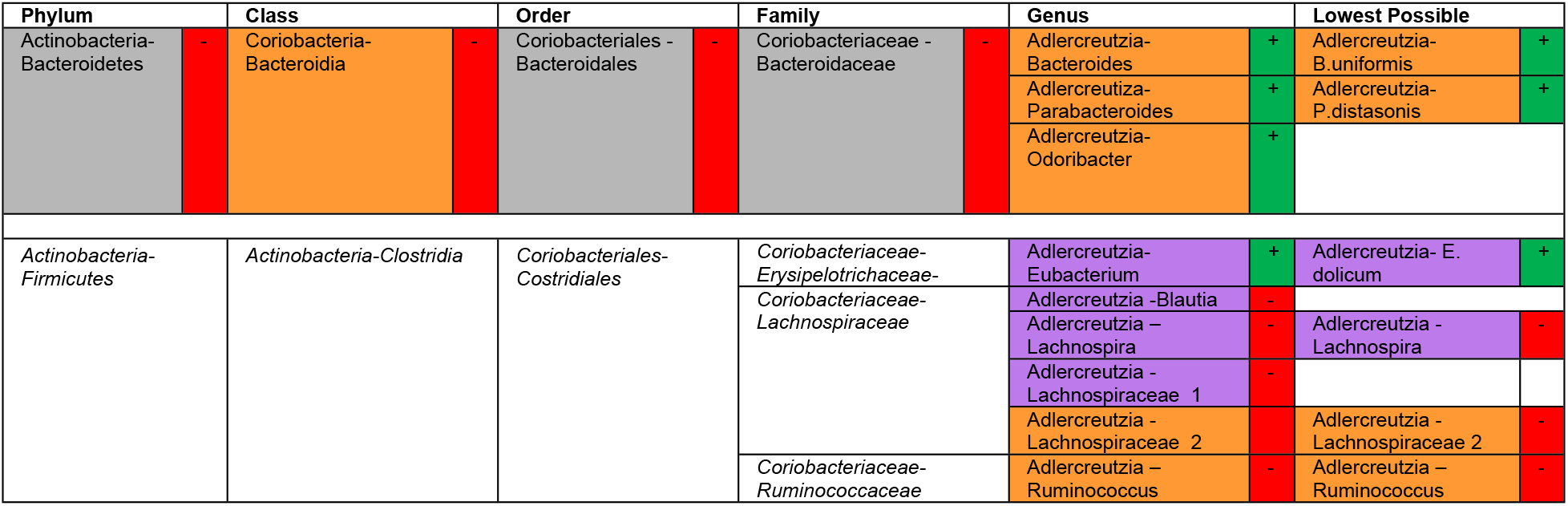
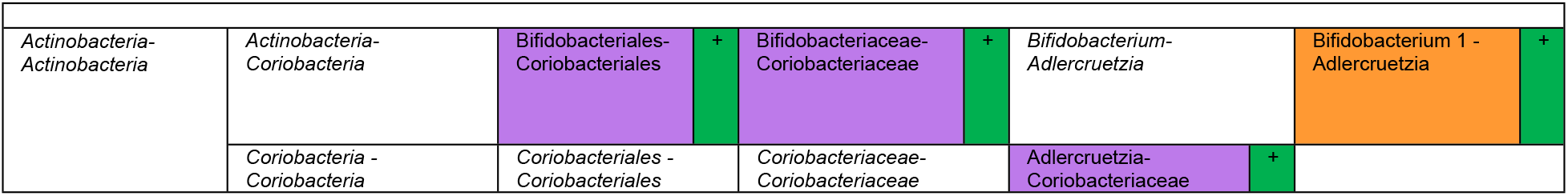
*Adlercruetzia* correlations.

**(F) Bacteroidetes-Firmicutes positive correlations (est. cooperation) are entirely exclusive to ADHD, and absent in Control.** Table 20 shows all Bacteroidetes-Firmicutes positive correlations. They are entirely limited to ADHD, and with one exception (*Clostridium*) involve Bacilli taxa.

**Table 20.**
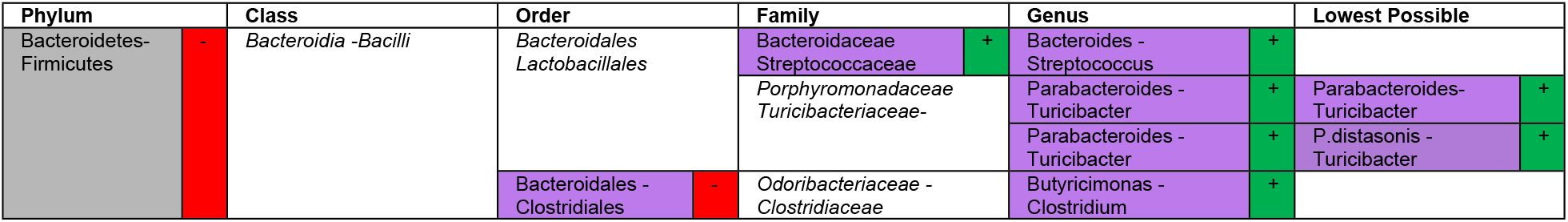
Bacteroidetes-Firmicutes positive correlations, over all MCNs.

We have already seen most of these, including *Clostridium* and ADHD LEfSe biomarker *Butyricimonas*, and the ADHD *FT*-*BB* connections involving *Turicibacter*, *Parabacteroides*, and *P.distasonis*. We now analyze the remaining top row, between *Bacteroides* (*Bacteroidaceae*) and *Streptococcus* (*Streptococcaceae*).

Firmicutes taxa were only ever present in cluster *BB* in ADHD, and we earlier noted *Streptococcaceae* and its genus *Streptococcus* as two of those taxa. Their cluster *BB* positive correlation was with centroid *Bacteroidaceae*/*Bacteroides*. Additionally cluster *BB* had almost no negative correlations (est. competition) with *FL*/*FR* (collectively 70% of the population) in ADHD, compared to a significant amount in Control.

What makes *Streptococcus* interesting for ADHD is that across all MCNs, it forms the only <u>positive</u> correlation between cluster *BB* and *FL/FR* (Fig. 7D). In other words, in addition to estimating significantly less *BB*-(*FL*/*FR*) competition in ADHD, our MCNs also estimate <u>cooperation</u> only in ADHD, between *Streptococcus* (*BB*) and *Blautia* (*FL1*).

Fig. 7D and 7F also show *Streptococcus* to be negatively correlated with *Oscillospira* in ADHD, a taxon we noted earlier its correlation sign change with *Enterobacteriaceae*. ATria (Table 21) also only ranks *Streptococcaceae*/*Streptococcus* as important in ADHD.

**Table 21.** ATria rankings of *Streptococcaceae*/*Streptococcus*.

| Phylum | Class | Order | Family | Genus | Lowest Possible |
| --- | --- | --- | --- | --- | --- |
| Firmicutes (#2/#T2) | Bacilli (NR/NR) | Lactobacillales (NR/NR) | Streptococcaceae (NR/#T12) | Streptococcus (NR/#T23) |  |

**(G) A shift in *Blautia*-*Oscillospira* dynamics, and their respective clusters.** Thus far *Oscillospira* has been noted for two ADHD-exclusive negative correlations, with taxa only ranked in ADHD: *Enterobacteriaceae* and *Streptococcus*. *Enterobacteriaceae*-*Oscillospira* was the only correlation to ever change sign from Control (positive) to ADHD (negative). *Streptococcus* was noted for its correlation with *Blautia*, the sole positive correlation between the largest Bacteroidetes-dominant cluster (*BB*) and Firmicutes-dominant clusters (*FL/FR*) in any MCN.

Previous studies have indicated butyrate-producing *Oscillospira* as a healthy gut taxon (118), specifically associated with leanness (119). *Blautia* is actually a taxon that has been associated with obesity (120). And interestingly in the Control MCN (Fig. 7C and 7E) *Blautia* and *Oscillospira* are negatively correlated, but not in ADHD (Fig. 7D and 7F).

Since obesity has been associated with ADHD (121), the shift in *Enterobacteriaceae* (*Oscillospira* cooperation in Control, competition in ADHD) and *Streptococcus* (*Blautia* cooperation and *Oscillospira* competition in ADHD) correlations become interesting, favoring *Blautia* cooperation and *Oscillospira* competition. Indeed correlation can never imply causation and further experimental verification is required. But ATria results (Table 22) also support this, ranking *Blautia* higher in ADHD and *Oscillospira* in Control.

**Table 22.** *Blautia* and *Oscillospira* ATria rankings (plus lineages). *=The lowest level had two *Blautia* taxa; we assumed the more abundant (*Blautia* 1, overall 9.3% relative abundance vs 0.6%, composing 93% of the *Blautia* population).

| Phylum | Class | Order | Family | Genus | Lowest Possible |
| --- | --- | --- | --- | --- | --- |
| Firmicutes (#T2/#2) | Clostridia (#T3/#T3) | Clostridiales (#T2/#3) | Lachnospiraceae (NR/NR) | Blautia (#T16/#13) | Blautia 1* (NR/#19) |
|  |  |  | Ruminococcaceae (#4/#T8) | Oscillospira (#3/#10) | Oscillospira (#5/#9) |

In fact our heatmap (Fig. 8C-8F) shows by intersecting *Oscillospira*’s row (small green rectangle) with the columns of *Blautia*’s cluster (gold rectangles, Control *FL3*, ADHD *FL1*) that Oscillospira is negatively correlated with *<u>Blautia</u>*<u>’s entire cluster</u> in Control, and these correlations are completely absent in ADHD.

The lowest level MCNs (Fig. 7E-F) also show *Blautia*’s cluster as larger in ADHD, and *Oscillospira*’s cluster as larger in Control. Table 23 contains members of these clusters. *Blautia* and *Oscillospira* each belong to a cluster dominated by its respective family: *Lachnospiraceae* (*FL*), and *Ruminococcaceae* (*FR*). *Oscillospira* is a core *FR* member and at the lowest level, we see the Control FR cluster (with *Enterobacteriaceae* now a member). *Blautia* is consistently a member of the same cluster as both *Lachnospiraceae* taxa in ADHD, comparably larger than its *FL3* Control cluster.

**Table 23.**
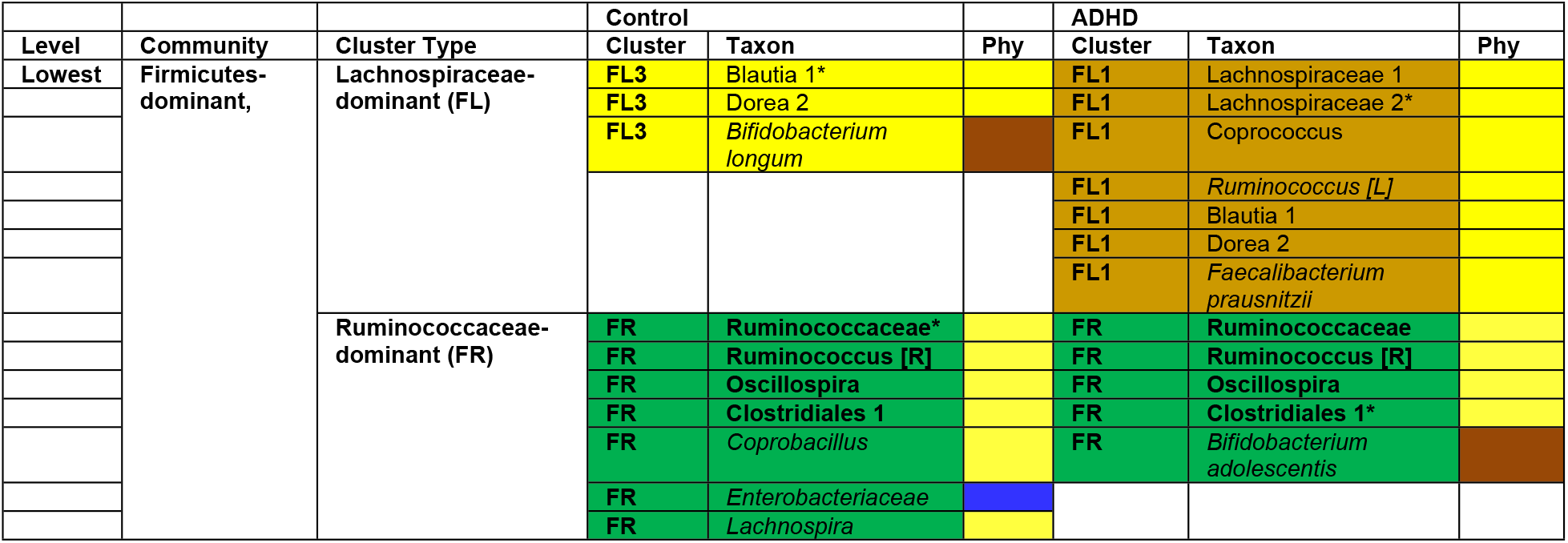
*Blautia* and *Oscillospira* clusters.

Heatmaps also indicate increased participation of *Oscillospira*’s cluster (*FR*) in Control (large green rectangle, Fig. 8C-F), including negative correlations with cluster *BB* that are absent in ADHD, yet another example of reduced ADHD cluster *BB* competition. In the MCNs, Fruchterman-Reingold places cluster *FR* (green) in a much more central position in Control (Fig. 7C vs. 7D, and 7E vs. 7F). The negative correlations between *Oscillospira* and *Blautia*’s entire cluster *FL3* (Fig. 7C and 7E) are also evident, almost separating FL3 from the MCN. In ADHD *Blautia*’s cluster *FL1* (gold) occupies a much more central position (Fig. 7D and 7F), with increased ADHD size particularly noticeable at the lowest level (Fig. 7F).

ATria (Table 24) indicates a general increased importance of *Blautia’s* family (*Lachnospiraceae*) in ADHD, and *Oscillospira’s* family (*Ruminococcaceae*) in Control. A couple of noteworthy taxa follow this trend. *Faecalibacterium prausnitzii (Ruminococcaceae)*, an anti-inflammatory bacterium (122) touted as a next-generation probiotic (123), is only ranked in Control. *Ruminococcus gnavis* (*Lachnospiraceae*), known to produce an inflammatory polysaccharide (124), is only ranked in ADHD.

**Table 24.** ATria rankings of *Lachnospiraceae* and *Ruminococcaceae* taxa.

| Phylum | Class | Order | Family | Genus | Lowest Possible |
| --- | --- | --- | --- | --- | --- |
| Firmicutes (#2/#T2) | Clostridia (#T3/#T3) | Clostridiales (#T2/#3) | *Lachnospiraceae* (NR/NR) | *Anaerostipes* (#T19/NR) | *Anaerostipes* (#6/#T26) |
|  |  |  |  | *Blautia* (#T16/#13) | *Blautia* 1 (NR/#19) |
|  |  |  |  |  | *Blautia* 2 (#9/#T22) |
|  |  |  |  | *Coprococcus* (NR/#5) | *Coprococcus* (#T21/#6) |
|  |  |  |  | *Dorea* (#T14/#12) | *Dorea* 2 (#13/NR) |
|  |  |  |  | *Lachnospira* (NR/#16) | *Lachnospira* (#T27/#5) |
|  |  |  |  | *Lachnospiraceae* 1 (#19/#15) | *Lachnospiraceae* 1 (#17/#2) |
|  |  |  |  | *Lachnospiraceae* 2 (#7/NR) | *Lachnospiraceae* 2 (#8/#14) |
|  |  |  |  | *Roseburia* (NR/NR) | *Roseburia* 1 (NR/#T28) |
|  |  |  |  |  | *Roseburia* 2 (#T25/#T22) |
|  |  |  |  | *Ruminococcus* (#1/#11) | *R. gnavis* (NR/#12) |
|  |  |  | *Ruminococcaceae* (#4/#T8) | *Faecalibacterium* (#T16/#2) | *F. prausnitzii* (#11/NR) |
|  |  |  |  | *Oscillospira* (#3/#10) | *Oscillospira* (#5/#9) |
|  |  |  |  | *Ruminococcaceae* (#8/#9) | *Ruminococcaceae* (#4/#18) |
|  |  |  |  | *Ruminococcus* (#2/NR) | *Ruminococcus* (#T21/NR) |

#### Summary

Four clusters were consistently present in both Control and ADHD MCNs. Three are Firmicutes-dominant (*FL*, *FR*, *FT*) and one is Bacteroidetes-dominant (*BB*). Table 25 shows their attributes, and summarizes observations we made about each.

**Table 25.** Largest, consistently present clusters.

| Cluster | Attribute | Observation |
| --- | --- | --- |
| BB | Largest Bacteroidetes-dominant cluster | Larger in ADHD, with more internal cooperation and less external competition. |
| FL(1,2,...) | Multiple Firmicutes, family *Lachnospiraceae*-dominant clusters | One large, centrally located cluster emerges in ADHD (FL1). Others are small, about the same size, and more disconnected (all are this way in Control). |
| FR | Firmicutes, family *Ruminococcaceae*-dominant | Smaller and less centrally located in ADHD. |
| FT | Firmicutes, core member *Turicibacter* | Slightly larger in ADHD. |

Table 26 summarizes correlations between members of these clusters. Other than the one exception in ADHD involving *Streptococcus* and *Blautia*: *FT* is the only Firmicutes-dominant cluster with taxa positively correlated with Bacteroidetes-dominant cluster (*BB)* members, and all correlations involving *FL*/*FR* (largest Firmicute-dominant clusters) and *BB* taxa are negative. *FT* is completely disconnected from *FL*/*FR* except some ADHD competition. *FL*-*FR* competition only happens in Control.

**Table 26.**
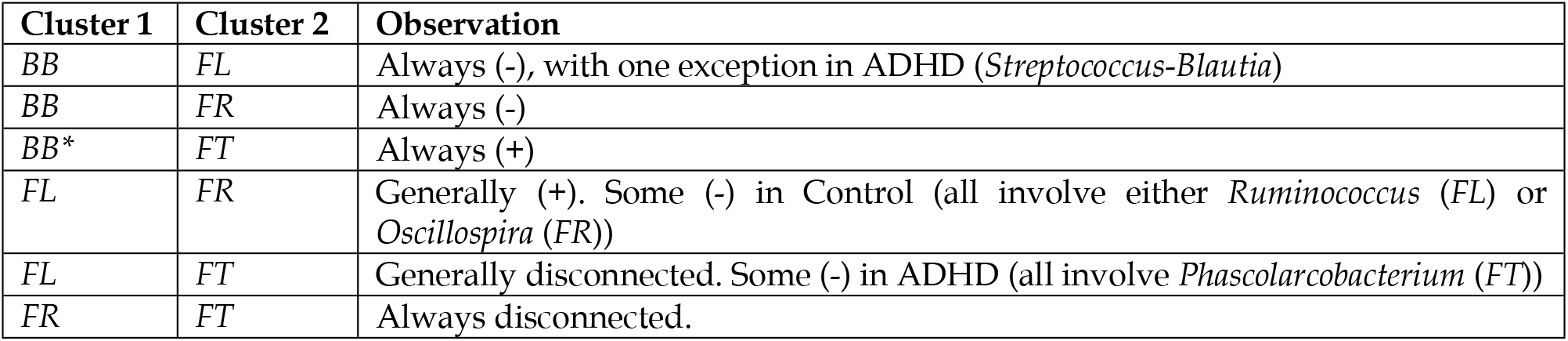
Interactions between taxa from Table 25 clusters (*=In Control, this took place with *BM1* after the *BB* “split”).

| Cluster 1 | Cluster 2 | Observation |
| --- | --- | --- |
| BB | FL | Always (-), with one exception in ADHD (*Streptococcus*-*Blautia*) |
| BB | FR | Always (-) |
| BB* | FT | Always (+) |
| FL | FR | Generally (+). Some (-) in Control (all involve either *Ruminococcus* (FL) or *Oscillospira* (FR)) |
| FL | FT | Generally disconnected. Some (-) in ADHD (all involve *Phascolarctobacterium* (FT)) |
| FR | FT | Always disconnected. |

Finally, we summarize taxa (Table 27) and relationships (Table 28) that we noted throughout our analyses.

**Table 27.**
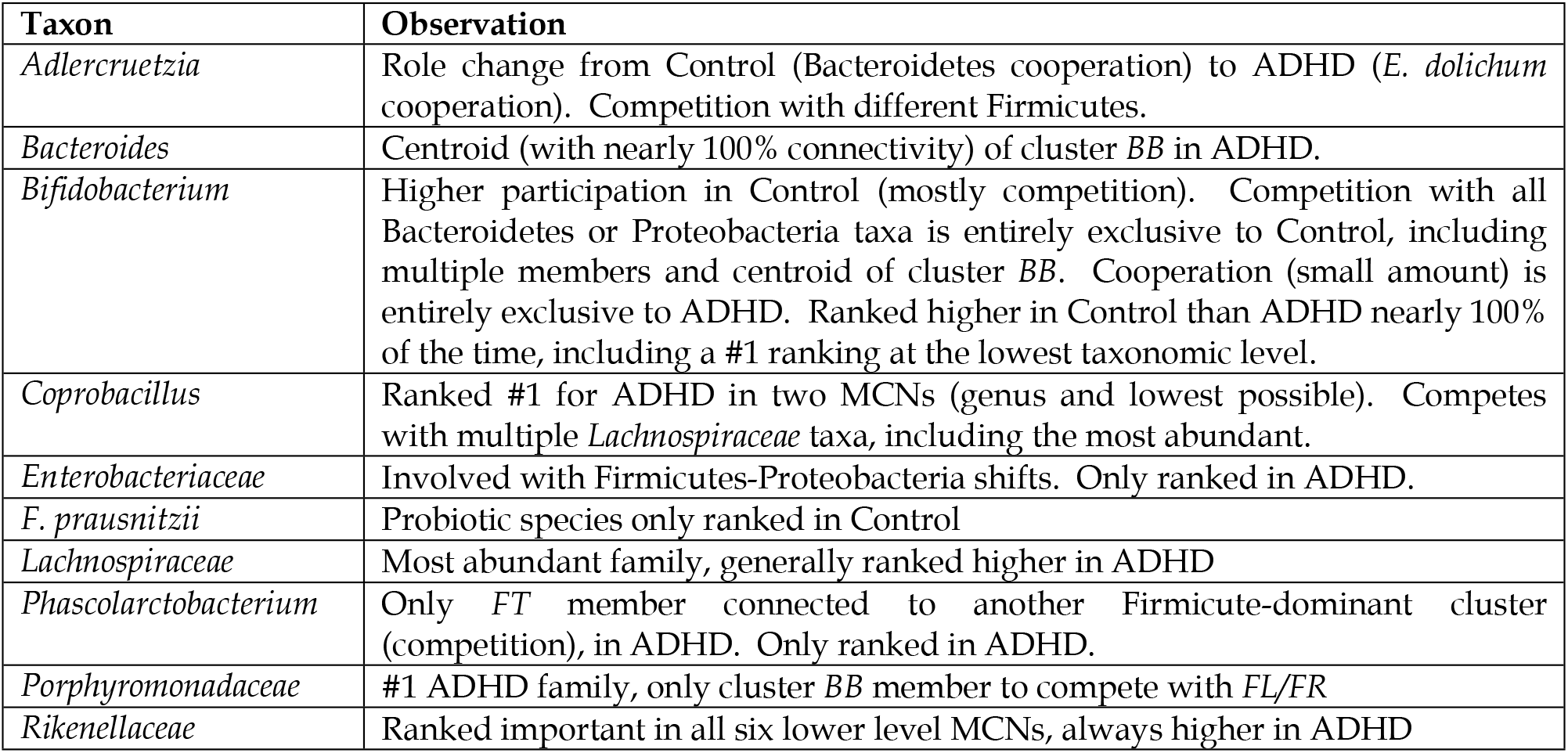
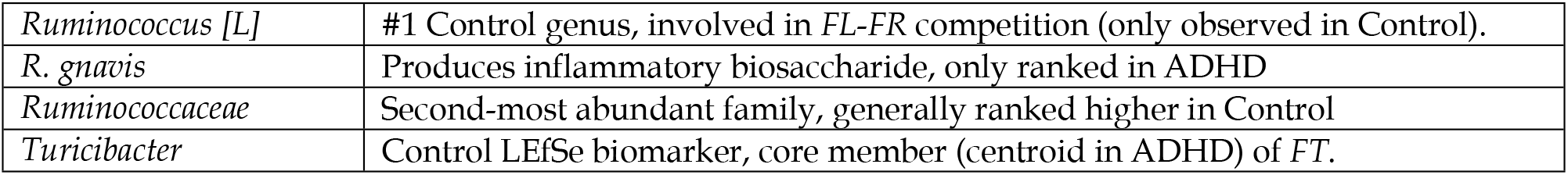
Taxa we noted throughout our analyses.

**Table 28.** Relationships noted throughout our analyses.

| Relationship(s) | Reason |
| --- | --- |
| *Bacteroides-Sutterella* (+, both) | Only core correlation consistent across both sample sets at all levels (12 MCNs). |
| *Bacteroides-Prevotella* (-, ADHD) | Only competition involving two Bacteroidetes taxa. |
| *Bacteroides-Streptococcus* (+, ADHD) | *Streptococcus* is one of only two Firmicutes genera to join cluster *BB*, through this connection. |
| *Butyricimonas-Clostridium* (+, ADHD) | *Butyricimonas* is an ADHD LEfSe biomarker, and only ranked by Atria in ADHD. *Clostridium* is one of only two Firmicutes to join cluster *BB*, through this connection. |
| *Clostridiaceae-Odoribacter* (-, ADHD) | *Odoribacter* is an ADHD LEfSe biomarker. In ADHD competes with *Clostridiaceae* taxa that compete with multiple cluster *BB* members (including its centroid). |
| *Enterobacteriaceae-Oscillospira* (+, Control; -, ADHD) | Only correlation ever to change sign from Control to ADHD. Taxa involved are in the same cluster in Control. |
| *Blautia-Oscillospira* (-, Control)<br>*Streptococcus-Blautia* (+, ADHD)<br>*Streptococcus-Oscillospira* (-, ADHD) | *Blautia* is associated with obesity and *Oscillospira* with leanness. *Oscillospira* (*FR*) competes with every member of *Blautia*'s cluster (*FL3*) in Control. *FL-FR* competition only happens in Control.<br>In ADHD *Streptococcus* cooperates with *Blautia* (obesity) and competes with *Oscillospira* (leanness). *Streptococcus-Blautia* is the only time a cluster *BB* member (largest Bacteroidetes-dominant) ever cooperates with taxa from *FL* or *FR* (largest Firmicutes-dominant, collectively over 70% of the population).<br>*Streptococcus* is only ranked in ADHD, *Blautia* is ranked higher in ADHD, *Oscillospira* is ranked higher in Control.<br>*Blautia*'s cluster (*FL1*) is larger and more central in ADHD. *Oscillospira*'s (*FR*) is larger and more central in Control. |

## Discussion

Traditional analysis methods (i.e. diversity and composition) prevalent in current ADHD gut microbiome literature provide a macroscale representation of a complex ecosystem. Conducting some of these approaches on equal-sized, gender-balanced undergraduate Control and ADHD gut microbiome datasets produced many results that corresponded with this literature, plus a potentially new Control biomarker *Turicibacter*. Current literature, as well as our results, suggest this macroscale perspective leaves a largely incomplete picture due to its neglect of underlying complexity. Our goal was to complete more of this picture by venturing deeper, by analyzing two-way ecological relationships (cooperation and competition), plus community detection, and centrality.

Our results provide a deeper meaning to those from the macroscale. Anomalous results involving elevated *Bifidobacterium* and reduced *Bacteroides* and *Sutterella* at ASRS extremes imposed significant challenges when interpreting results (with *Bifidobacterium*, we were not the first to observe this (58)). Our MCNs estimate that a Bacteroidetes-dominant community (cluster *BB*) forms in both microbiomes, with *Bacteroides* and *Sutterella* both core members, that in ADHD is larger, more centered around *Bacteroides*, residing in conditions that favor its <u>cooperation</u>, as opposed to <u>competition</u> in Control. And our MCNs estimate *Bifidobacterium* to be involved in these conditions, shifting from exclusively <u>competitive</u> relationships with cluster *BB* members (including its most abundant and centroid) in Control, to exclusively a <u>cooperative</u> relationship in ADHD.

Potential roles played by LEfSe biomarkers also became observable. Our MCNs estimated *Odoribacter*, reported by our LEfSe analysis and another (59) as ADHD-elevated, to also compete with two *Clostridiaceae* taxa that competed with cluster *BB* taxa. Another one of our ADHD biomarkers, *Butyricimonas*, joined cluster *BB* in ADHD and formed cooperative relationships with many members. New interesting taxa and communities also emerged. Cluster *FT* (cooperative with cluster *BB*) was larger in ADHD. Cluster *FR* (*Ruminococcaceae*-dominant, competitive with cluster *BB*) was smaller in ADHD, with *Ruminococcaceae* taxa almost universally less central. *Ruminococcaceae* member genus *Oscillospira* was estimated to have ADHD-exclusive competition, with *Enterobacteriaceae* (cooperative and fellow *FR* member in Control), and cluster *BB* member *Streptococcus*. The shift in dynamics from Control to ADHD involving *Streptococcus*, *Blautia*, and *Oscillospira* in ADHD was particularly interesting (Table 28, last row).

Deeper meaning can be added through additional studies targeting some of these taxa and relationships, including multi-omics (125) and/or physical laboratory experiments. Fundamentally, ecological relationships manifest through internal interplay within the underlying web of interactions (126). Cooperation could take place for example if two taxa produce a nutrient that the other consumes; competition could take place if two taxa consume a nutrient that neither produces. Coupling taxa to metabolites they produce and consume and analyzing pathways can help elucidate underlying mechanisms behind these ecological relationships. These pathways can then be searched for neurotransmitters to establish ADHD connections. With very few studies even attempting this level of analysis (62), an enormous breadth of knowledge remains.

Many future improvements to our analyses are possible. Future studies involving ADHD and the gut microbiome should account for factors such as ethnicity (127), use of medication/probiotics (55), use of antibiotics (128), diet (129), and gastrointestinal issues (130). More meaning to relationships in our MCNs can also be uncovered, through *causality* studies. Causality would give direction to edges, enabling detection of both two- and one-way (i.e. commensalism (110), amensalism (111)) relationships. This can be achieved through for example Bayesian Networks (131), which detect relationships where a taxon is conditionally dependent on another. Conditional dependence also eliminates spurious edges that can occur with correlations; for example, two entities that co-occur with a mutual entity will naturally tend to co-occur (88) (this was also a dependency removed by ATria after finding a central node). Sazal *et al*. (132) have already verified such networks as a predictor for oral microbiome colonization order. Time can also factor into ecological relationships because while sometimes these relationships are constant in microbiomes (133), they can also be transitive (134) or even time-varying (135). DBNs that account for time have already been used to predict long-term infant gut behavior (136). Higher-level network metrics such as modularity (137) and vulnerability (138) would provide another potential avenue for comparing and contrasting Control and ADHD MCNs. Amplicon Sequence Variants (ASVs, (139)) can be used in place of the current Operational Taxonomic Units (OTUs) that are generated by similarity-based clustering. ASVs exhibit more reliability at lower levels of the taxonomic tree and can improve the granularity of our MCNs, achieving more species- and sometimes even strain-level classifications.

A more complete understanding of ADHD and the gut microbiome will best equip the community to make the right decisions when administering treatment(s). Our results, coupled with those in the literature, suggest that the gut microbiota cannot afford to be ignored when it comes to ADHD, and treatments directly targeting the gut microbiome have potential. Encouraging results have been uncovered for gluten and casein-free diets (44), Microbiota Transfer Therapy (MTT, (140,141)), and probiotics (142) with ASD. Our results also indicate that the gut microbiome is an ecosystem, and any changes to one single element will likely impact other members. Additionally since the human gut microbiome is widely varied across individuals (143), personalized medicine should be used when developing such treatments.

## Acknowledgements

The authors would like to thank colleagues from the Bioinformatics Research Group (BioRG) and the Stollstorff Lab for many useful discussions. The work of GN and KM was supported by the National Institute of Health [Award #580 1R15AI128714-01]. The work of GN was also supported by the Department of Defense [Contract #581 W911NF-16-1-0494] and National Institute of Justice [Award #582 2017-NE-BX-0001]. TC received support from NVIDIA and Florida International University.

## Supporting Information Captions

**Fig. S1.**
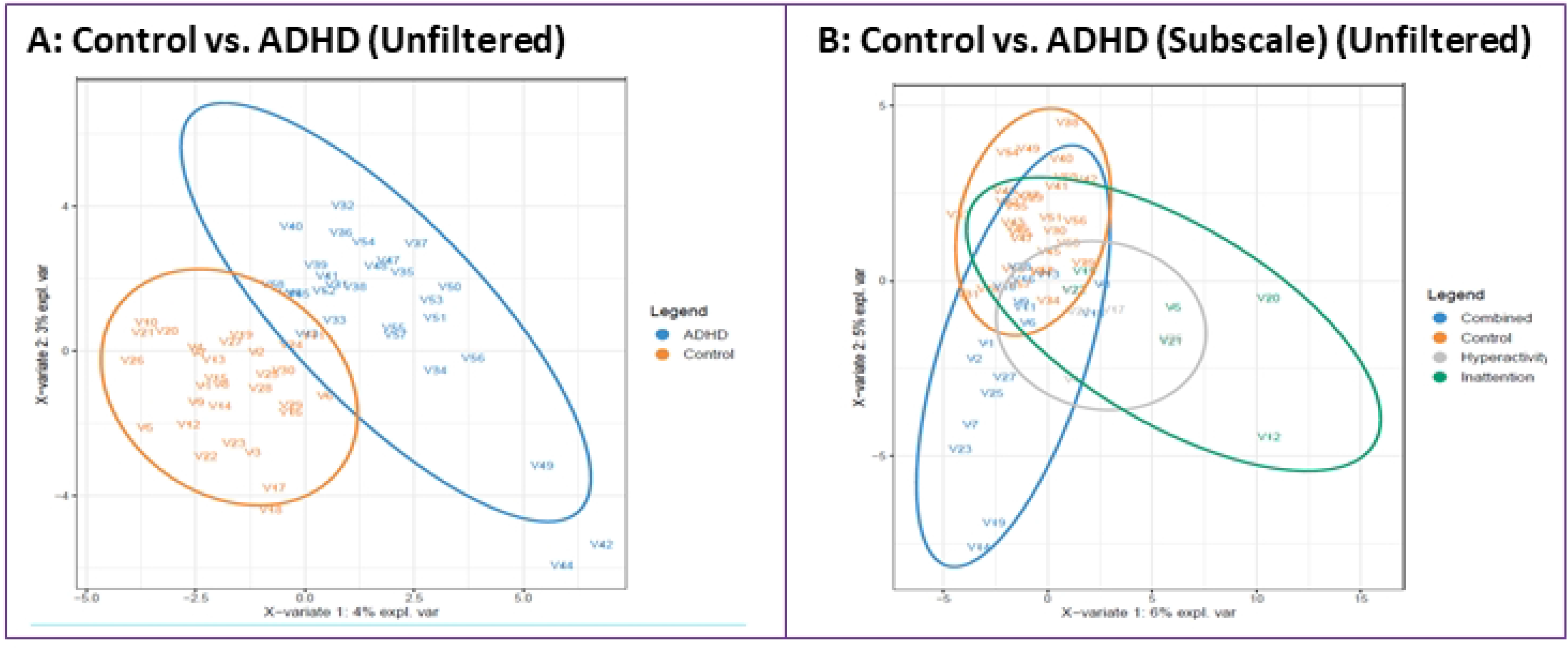
Discriminant Analysis, Scarce Taxa Included. Results of running sPLS-DA (68) on microbiome abundance data (ellipse confidence level 95%) without removing scarce taxa. The figures show the analyses (a) comparing Control (orange) and ADHD (blue) groups and (b) further separating the ADHD group into inattention (green), hyperactive (grey), and combined (blue).

**Fig. S2.**
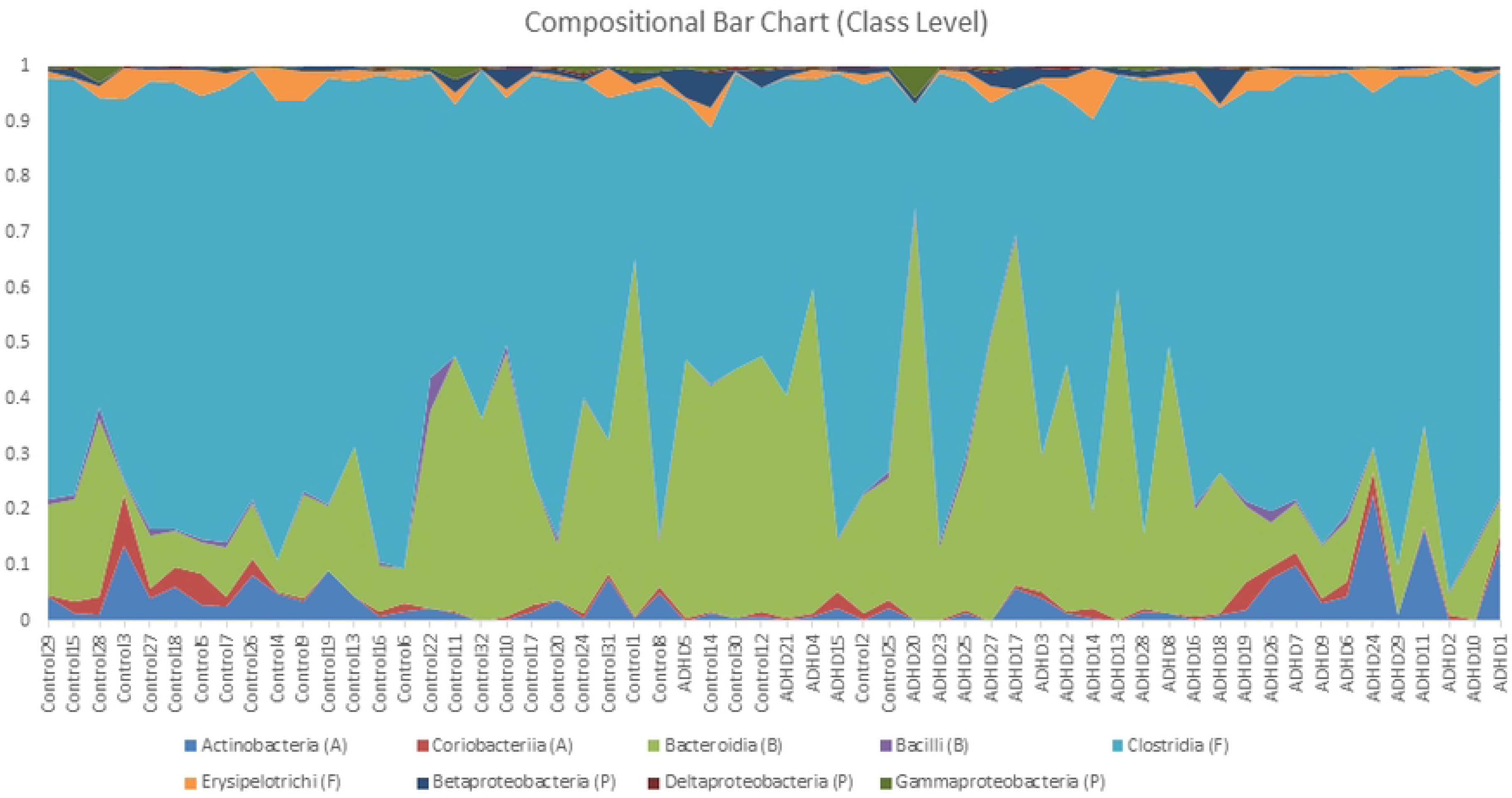
Compositional Analysis, Class Level. Microbial compositional bar graph for each subject, generated using QIIME (70), conducted at the class level. Subjects are ordered by increasing Adult ADHD Self Report Scale (ASRS) score, with the y-axis representing relative abundance.

**Fig. S3.**
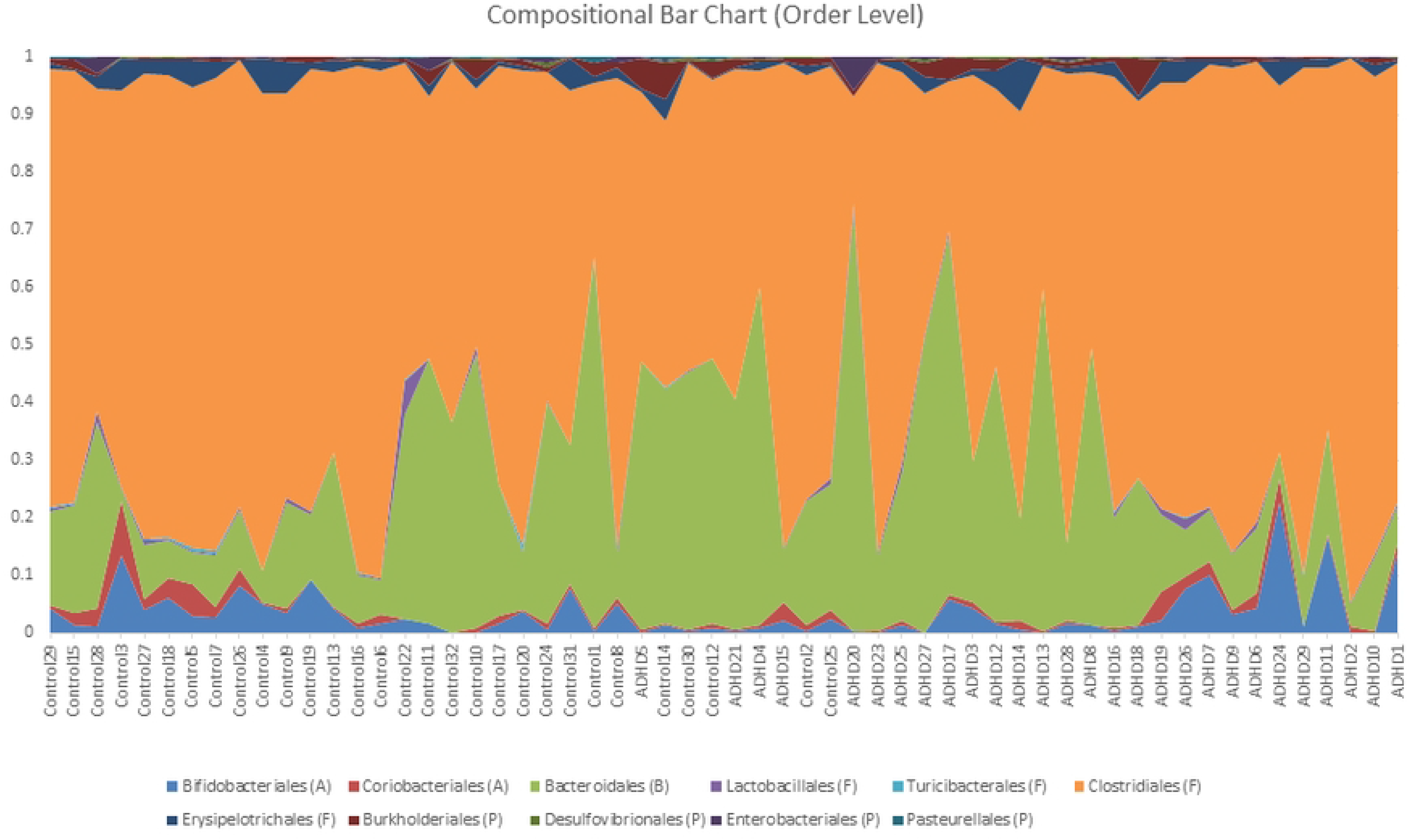
Compositional Analysis, Order Level. Microbial compositional bar graph for each subject, generated using QIIME (70), conducted at the order level. Subjects are ordered by increasing Adult ADHD Self Report Scale (ASRS) score, with the y-axis representing relative abundance.

**Fig. S4.**
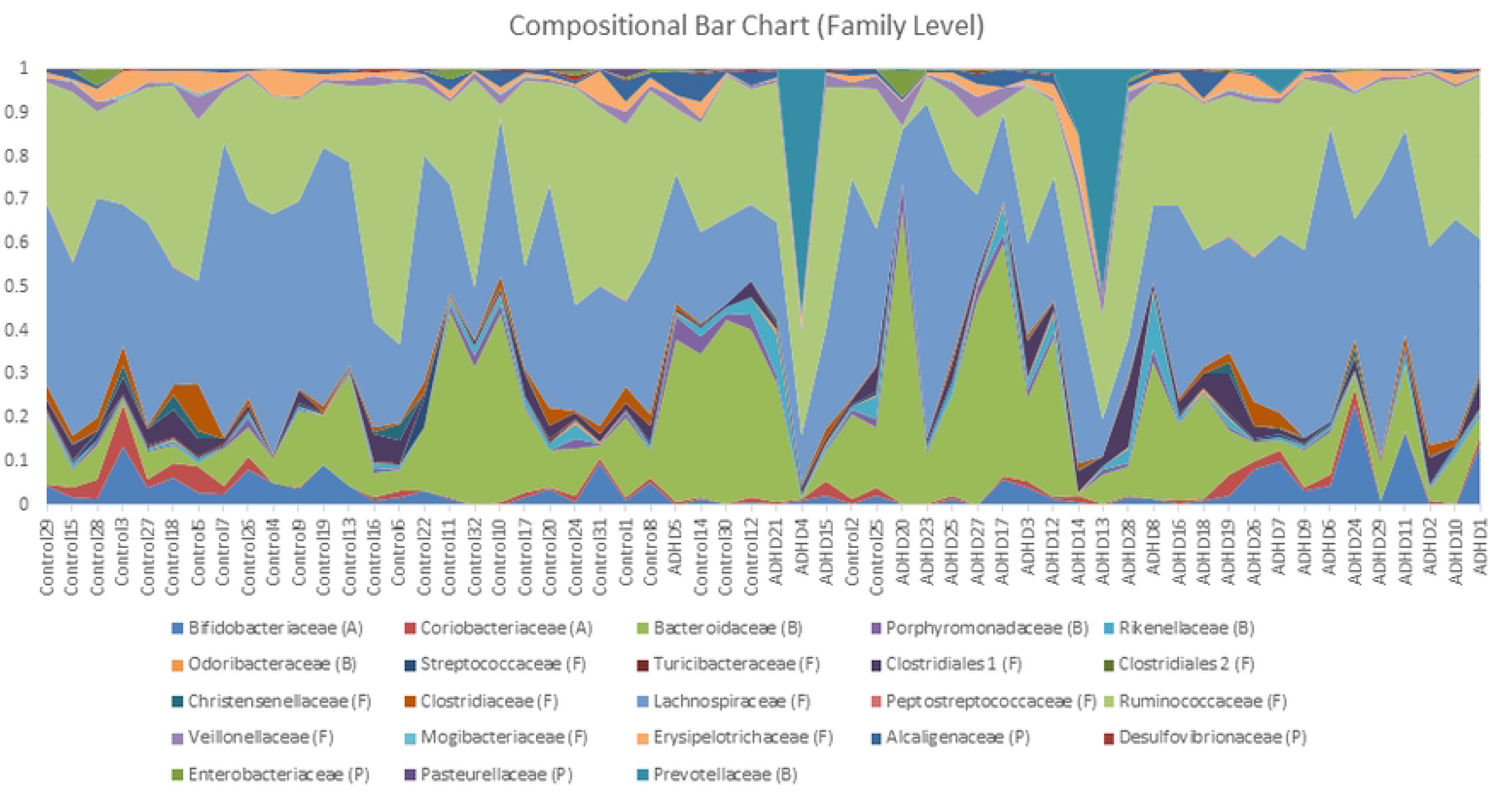
Compositional Analysis, Family Level. Microbial compositional bar graph for each subject, generated using QIIME (70), conducted at the family level. Subjects are ordered by increasing Adult ADHD Self Report Scale (ASRS) score, with the y-axis representing relative abundance.

**Fig. S5.**
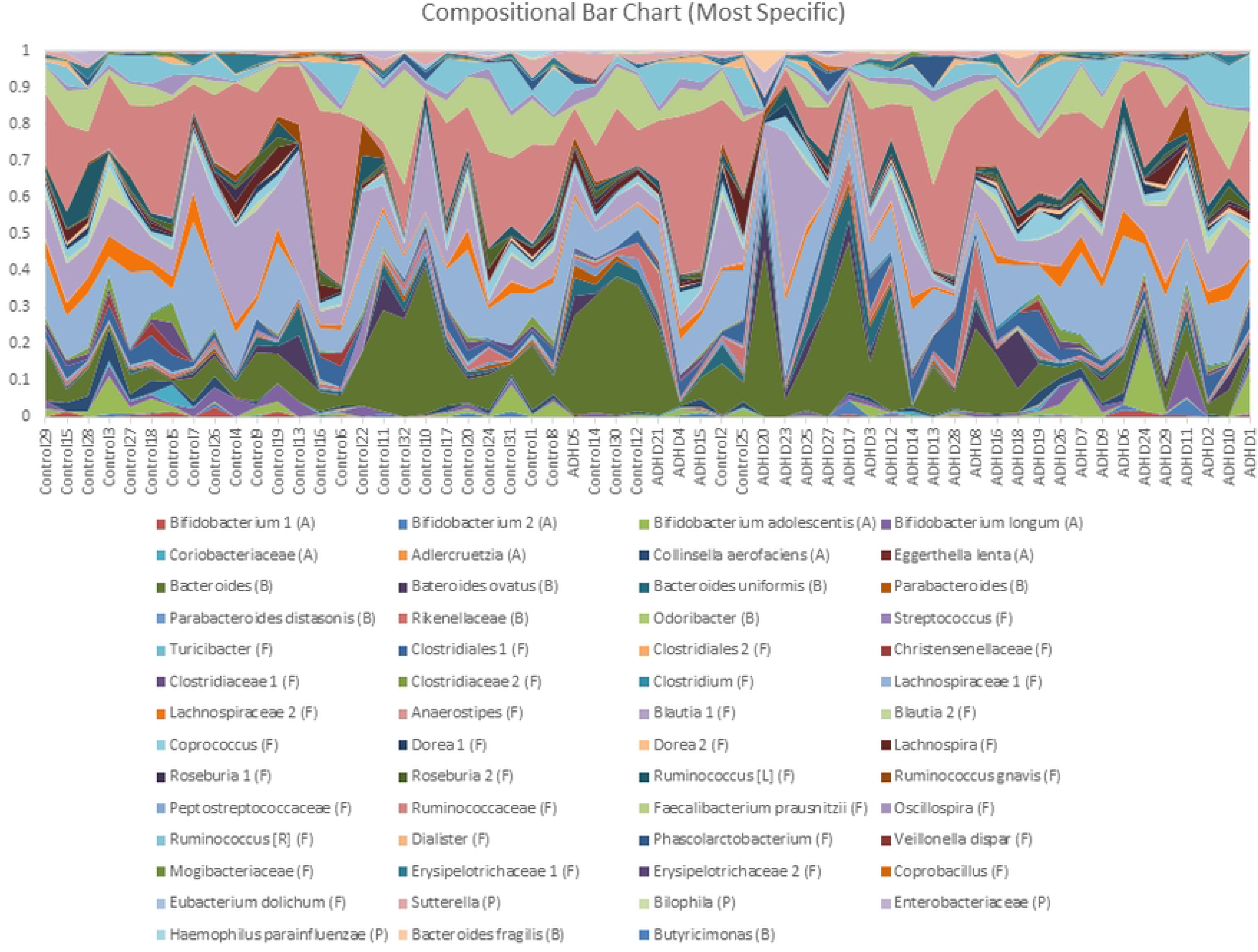
Compositional Analysis, Order Level. Microbial compositional bar graph for each subject, generated using QIIME (70), conducted at the species level. Subjects are ordered by increasing Adult ADHD Self Report Scale (ASRS) score, with the y-axis representing relative abundance.

**Fig. S6.**
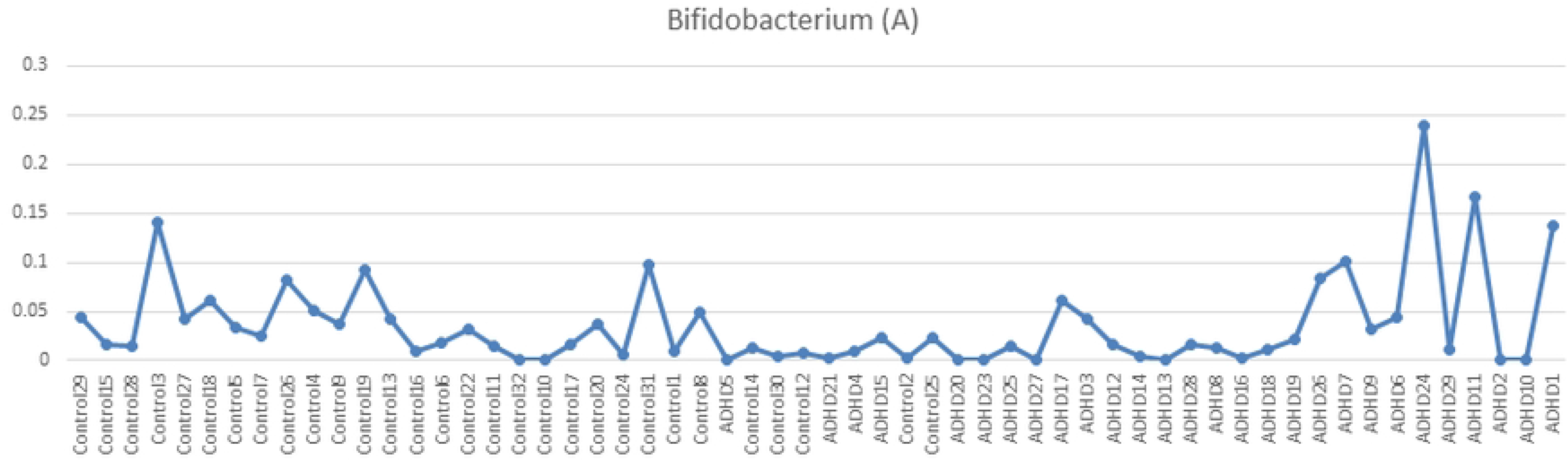
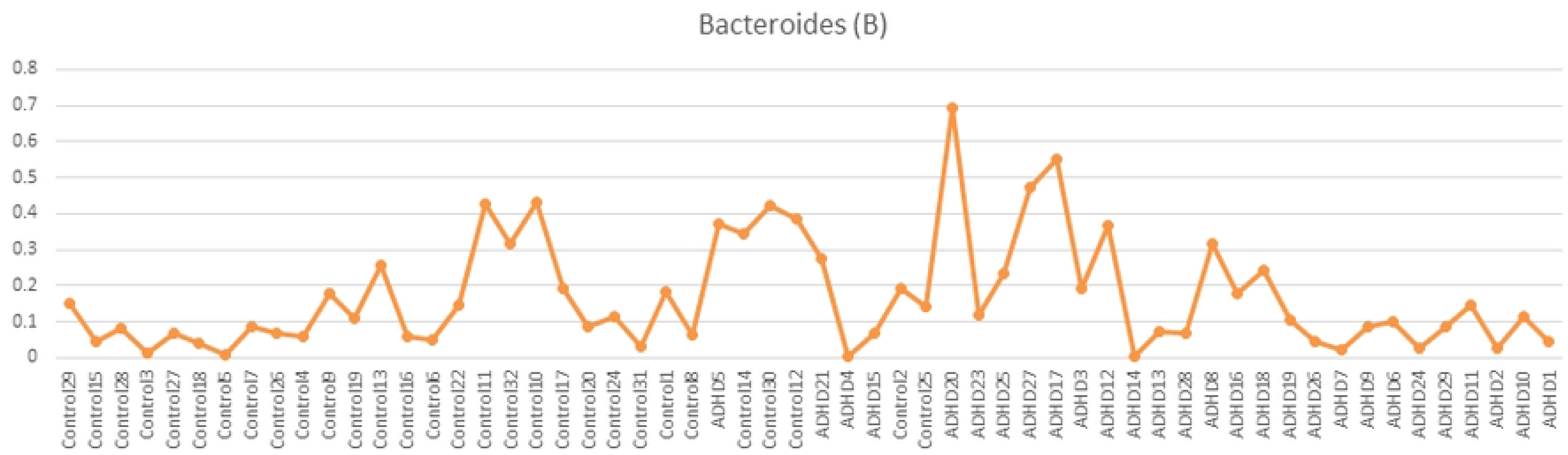
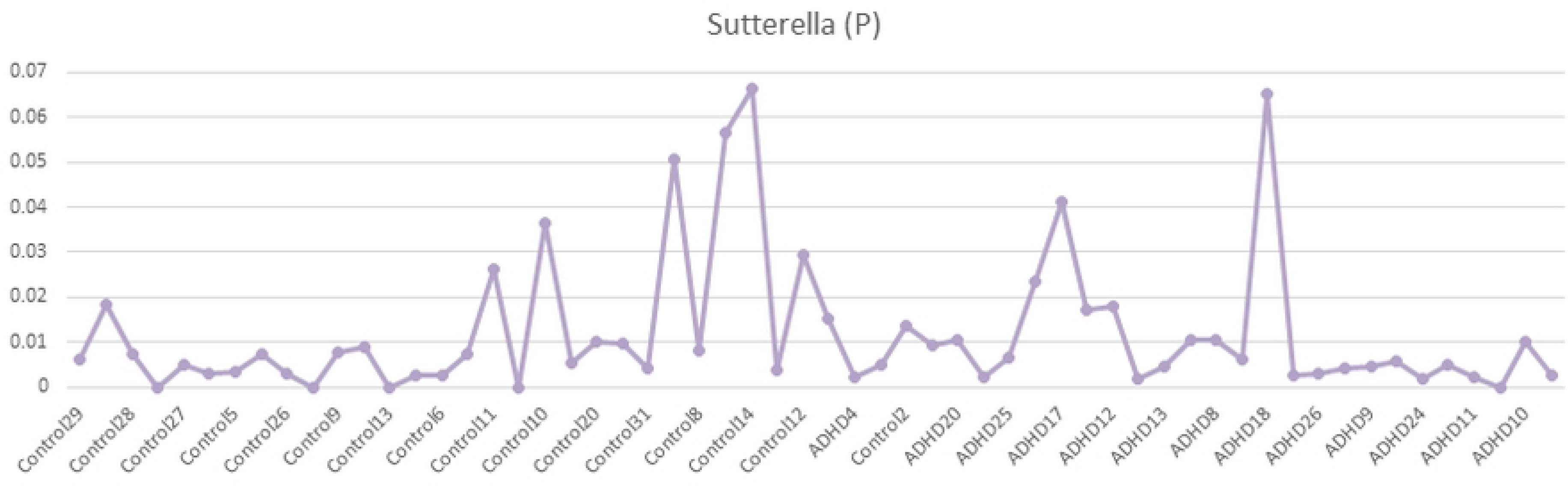
Relative Abundance, Three Observed Taxa. Relative abundance of genera (a) *Bifidobacterium*, (b) *Bacteroides* and (c) *Sutterella*. Subjects are ordered by increasing Adult ADHD Self Report Scale (ASRS) score, with the y-axis representing relative abundance.

**Table S1.** Correlations in all MCNs, over all taxonomic levels, organized by taxonomic classification. Each box indicates the two taxa involved in each correlation, along with the sign (+ or -). Boxes colored orange correspond to correlations on present in Control, and purple only present in ADHD. Grey boxes are present in both MCNs. White boxes correspond to correlations that were not observed, but one was present among its descendants (i.e. genera *Collinsella* and *Butyricimonas* were not correlated in either MCN, but member taxa *C. aerofaciens* and *Butyricimonas* were for ADHD). For polyphetic genus *Ruminococcus*, [L]=*Lachnospiraceae* family, [R]=*Ruminococcaceae* family.

**Table S2.** ATria rankings of all taxa found as important in all MCNs, grouped by taxonomic classification. NR=Not Ranked, T=Tied. Taxa ranked only in Control are colored dark orange, higher in Control light orange, higher in ADHD light purple, and only in ADHD dark purple. **Bold** taxa are ranked #1 in their corresponding MCN. White, italicized taxa correspond to unranked taxa with a ranked descendant.

**Table S3.** Family-level MCN clusters, reported by Affinity Propagation (AP, [CITE]). Core taxa (shared by both MCNs) are bold, and centroids are marked with an asterisk (*, requires at least three taxa). Italicized taxa are exclusive to their MCN (Control or ADHD). Phylum colors match those in Fig. 5 (Actinobacteria brown, Firmicutes yellow, Proteobacteria blue, Bacteroidetes dark purple). Cluster colors match those in Fig. 6.

**Table S4.** Genus-level clusters, reported by AP. Color and labelling is the same as Table S3. [L]=*Lachnospiraceae* family, [R]=*Ruminococcaceae* family.

**Table S5.** Lowest-level clusters, reported by AP. Color and labelling is the same as Tables S3 and S4.

## Notes

### Competing Interest Statement

The authors have declared no competing interest.

